# Pathogen-driven reactivation of metabolite prodrugs defines nitroxoline’s iron-deprivation antibiotic activity

**DOI:** 10.64898/2026.06.11.731531

**Authors:** Felix Deschner, Thorsten Kinsinger, Alexander F. Kiefer, Jürgen Bartel, Alexander Voltz, Kai Schließmann, Nina Walzer, Sören L. Becker, Andreas M. Kany, Franziska Fries, Jennifer Herrmann, Rolf Müller

## Abstract

*The rise of antimicrobial resistance warrants renewed attention to established but overlooked antibiotics such as nitroxoline (NTX). Here, we systematically dissect NTX’s mode of action and investigate the contribution of its first-pass metabolites, NTX-sulphate and NTX-glucuronide. We identified metallophore-mediated cellular iron deprivation as the principal antibacterial mechanism of NTX, characterized by induction of iron acquisition pathways and Fe–S cluster proteins, and concomitant loss of protein-bound iron. In contrast, NTX metabolites were biologically inactive and lacked metal-chelating properties. Ex vivo assays demonstrated that clinically relevant uropathogens, including Escherichia coli and Klebsiella pneumoniae, efficiently reconvert these metabolites into active NTX in human urine. Together, our findings establish a mechanistic framework linking NTX antibacterial activity, host detoxification, and pathogen-dependent metabolite reactivation, and providing a molecular explanation for NTX’s enduring therapeutic potential and favourable safety profile*.

## Introduction

Nitroxoline (5-nitro-8-hydroxyquinoline, NTX) is an antibiotic approved for the first-line treatment of uncomplicated urinary tract infections (uUTIs) caused *e.g.*, by *Escherichia coli* or *Klebsiella pneumoniae*. Beyond these indications, NTX exhibits a broad-spectrum antimicrobial activity (Deschner *et al*, 2024; Papareddy *et al*, 2025). With more than 50 years of clinical use, NTX is regarded as a long-established antimicrobial agent with a well-documented safety and efficacy profile. Despite its proven effectiveness, patient safety, and lack of resistance development, NTX is largely underused and has been almost forgotten by pharmaceutical industry and physicians (Deschner *et al*, 2024; Schmiemann *et al*, 2018; Kresken & Körber-Irrgang, 2014; Wagenlehner *et al*, 2014; Plambeck *et al*, 2022; Stoltidis-Claus *et al*, 2023). The emergence of multidrug-resistant human pathogens and the global spread of antimicrobial resistance (AMR) posing a threat to mankind (Global action plan on antimicrobial resistance, 2015; Murray *et al*, 2022), have brought NTX back into the focus of research. But surprisingly, the precise mode of action (MoA) of NTX and especially the activity of its human metabolites remained elusive. It is known that after oral application NTX is rapidly and almost completely metabolised to NTX-sulphate (NTX-S) and NTX-glucuronide (NTX-G) *via* phase-II metabolism prior to excretion into the urine (Wagenlehner *et al*, 2014). Therefore, although NTX is excreted mainly as NTX-S and NTX-G, the high efficacy of NTX in uUTI raises questions about the role and fate of these metabolites in mediating the biological effect.

Several MoA hypotheses have been proposed so far, mainly based on the previously reported chelator activity of NTX towards divalent cations. NTX was shown to directly coordinate, *e.g.*, Zn^2+^ or Cu^2+^ as a bidentate ligand resulting in the enzymatic inhibition of RNA polymerase or the methionine aminopeptidase by cofactor deprivation, as well as leading to reduced biofilm formation and adhesion capabilities to bladder epithelium, or effects of interference with bacterial membranes (Papareddy *et al*, 2025; Wagenlehner *et al*, 2014; Pelletier *et al*, 1995; Hu *et al*, 2013; Fraser & Creanor, 1974; Bourlioux *et al*, 1989; Sobke *et al*, 2012; Shim *et al*, 2010). Recently, NTX was shown to act as a metallophore in *Acinetobacter baumannii*, facilitating the intracellular accumulation of Zn²⁺ and Cu²⁺ and thereby inducing toxic metal overload. This intracellular excess of transition metals disrupts metal homeostasis and compromises iron–sulfur cluster proteins, ultimately affecting bacterial iron metabolism (Cacace *et al*, 2025). Surprisingly, NTX was also described as highly cytotoxic *in vitro* and as a candidate for anti-cancer therapy (Papareddy *et al*, 2025). It was further shown to be effective against brain-eating amoeba (Spottiswoode *et al*, 2023), as well as mpox virus (Bojkova *et al*, 2023). Unlike its antibacterial activity, the antiviral effect was not affected by the addition of divalent cations, suggesting that the underlying antiviral and antibacterial MoAs are different.

To better understand the antibacterial activity of NTX, we revisited its MoA in the model organism *E. coli* using an integrated approach combining global proteomics and size-exclusion chromatography coupled to elemental analysis (SEC-ICP-MS). We further chemically synthesised the major NTX metabolites, NTX-S and NTX-G, and showed them to be devoid of intrinsic antibacterial activity. Based on these findings, we developed a hypothesis for NTX activity during infection and demonstrate re-activation of these metabolites through pathogen-dependent bacterial biotransformation in human urine.

## Results

### Nitroxoline binds divalent metal ions with high affinity

NTX is a known chelator of divalent cations (**Figure SI 1**) (Lipunova *et al*, 2014), yet its metal-binding interactions have not previously been quantitatively characterised. We performed isothermal titration calorimetry using NTX and a variety of metal cations as chloride salts to gain insights into the general binding capabilities of NTX and to identify potential non-binding ligands.

We compared the monovalent cations sodium and potassium, the divalent main-group cations magnesium and calcium, and the transition metals zinc, iron, cobalt, manganese, nickel and copper, and for the first time quantitatively characterised NTX–metal interactions by determining dissociation constants (K_D_), enthalpy (dH), free energy (dG) and binding stoichiometry (N) at pH 6.2, mimicking conditions during uUTIs (**Figure 1**A/B, **Figure SI 2** – **Figure SI 10**). We could confirm binding of NTX to Zn^2+^, Co^2+^, Mn^2+^, Fe^2+^, Ni^2+^, and Cu^2+^, but neither Mg^2+^ or Ca^2+^, nor the monovalent ions K^+^ and Na^+^ showed any binding. While earlier studies emphasized an influence of Mg²⁺ and Mn²⁺ on NTX activity (Pelletier *et al*, 1995), our direct binding data are more consistent with recent MoA hypotheses describing NTX as a metallophore with preference for transition metal cations (Cacace *et al*, 2025).

**Figure 1:**
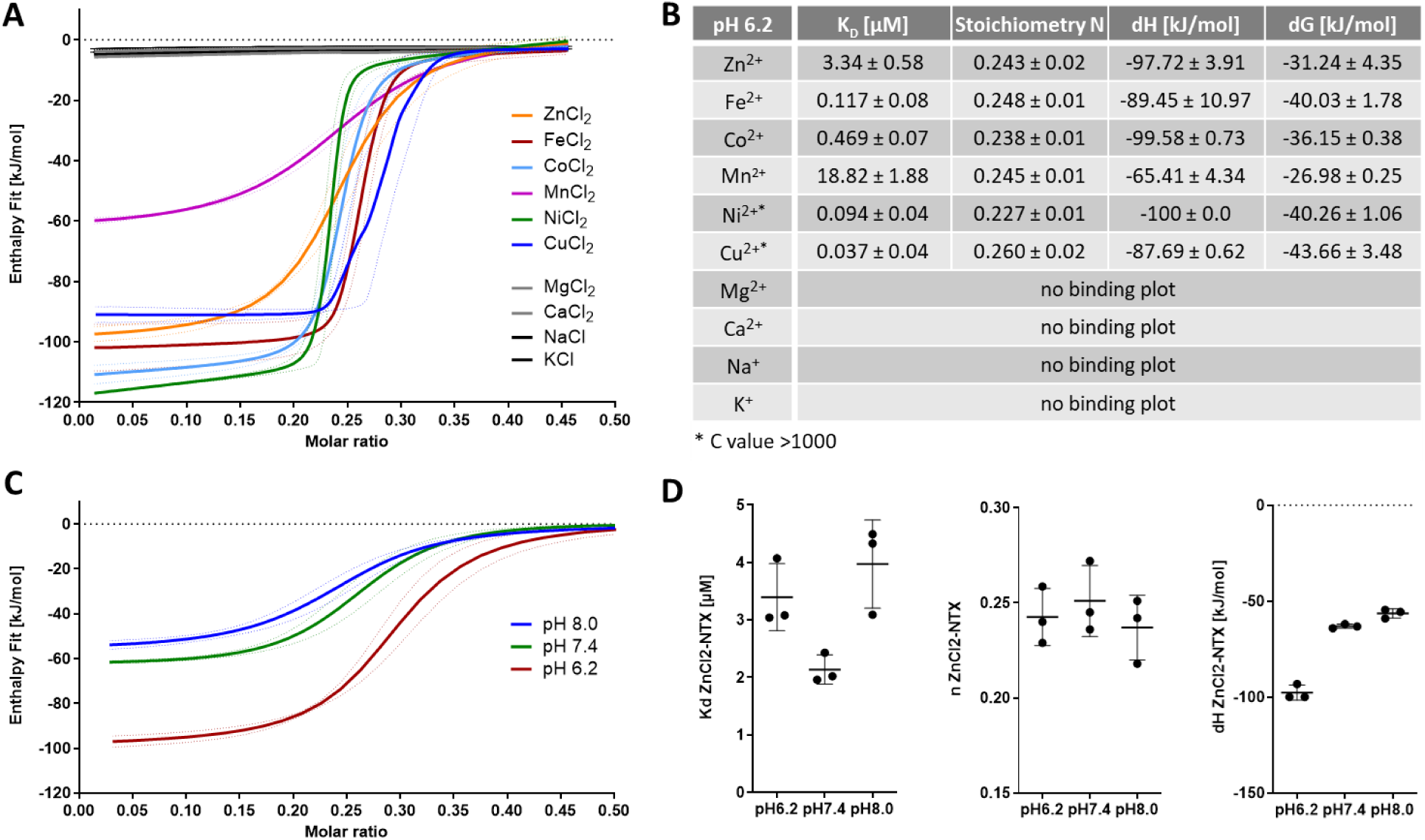
Isothermal titration calorimetry (ITC) confirms exothermic binding of NTX to divalent cations. A) Metal-to-ligand titration experiments were performed in triplicates in 100 mM Tris-HCl at pH 6.2 and 10% DMSO, with NTX in the cell and chloride salt in Tris-HCl in the syringe (for exact ratios, see **Figure SI 2**-**Figure SI 10**). Mean and SD are shown. B) Quantitative evaluation of ITC experiments using both the independent model and the blank (linear) model to adjust for base line shifts. C) pH dependent binding plot of NTX:Zn in 100 mM Tris-HCl at pH 6.2 (red), 7.4 (green) or 8 (blue). Mean and SD is shown (n = 3). D) Quantitative evaluation of pH-dependent Zn-NTX-complex formation.

In contrast to reported crystal structures where two NTX molecule coordinate one metal ion ([NTX_2_M]), reaction stoichiometry in ITC was repeatedly determined as 1:4 (ion:NTX), indicating π-stacking of NTX molecules or further effects of transition metal states in this experimental set up (Hu *et al*, 2013; Lipunova *et al*, 2014; Aziz *et al*, 2025). Overall, we found the K_D_ decreasing in the order: Mn^2+^ > Zn^2+^ > Co^2+^ > Fe^2+^ > Ni^2+^ > Cu^2+^, with the complex formation being exothermic and favourable as indicated by negative reaction enthalpy dH and free energy dG. Reaction c values were above 1000 for Ni^2+^ and Cu^2+^, rendering K_D_ determination unreliable. For Zn^2+^, we representatively showed NTX-binding at all physiological/urine relevant pH conditions (6.2, 7.4 and 8.0), and confirmed higher reaction enthalpy at acidic, rather than neutral or alkaline conditions (**Figure 1**C/D).

### NTX induces iron deficiency in *E. coli*

To understand the cellular response to NTX stress, we treated *E. coli* ATCC25922 with NTX and performed both, a full proteome analysis and SEC-ICP-MS for size-fractionated intracellular metal analysis. Cells were exposed to sub-inhibitory concentrations of NTX (0.25-fold MIC, 1 µg/mL; *n* = 4) and grown to early stationary phase prior sample preparation.

In full proteome experiments, more than 2,000 proteins have been identified confirming excellent protein coverage with 53 and 47 proteins significantly up-/or downregulated (log_2_ > 1.5 or log_2_ < -2.5), respectively. Different thresholds have been used to keep the number of proteins analysed in a reasonable range and to avoid false positive clustering. For evaluation, data were analysed using GO term (biological function) analyses via String-Networks (string-db.org) (**Figure 2**A/B) as well as KEGG pathway analysis using proteomaps (proteomaps.net, **Figure SI 11**, **Table SI 1**).

**Figure 2:**
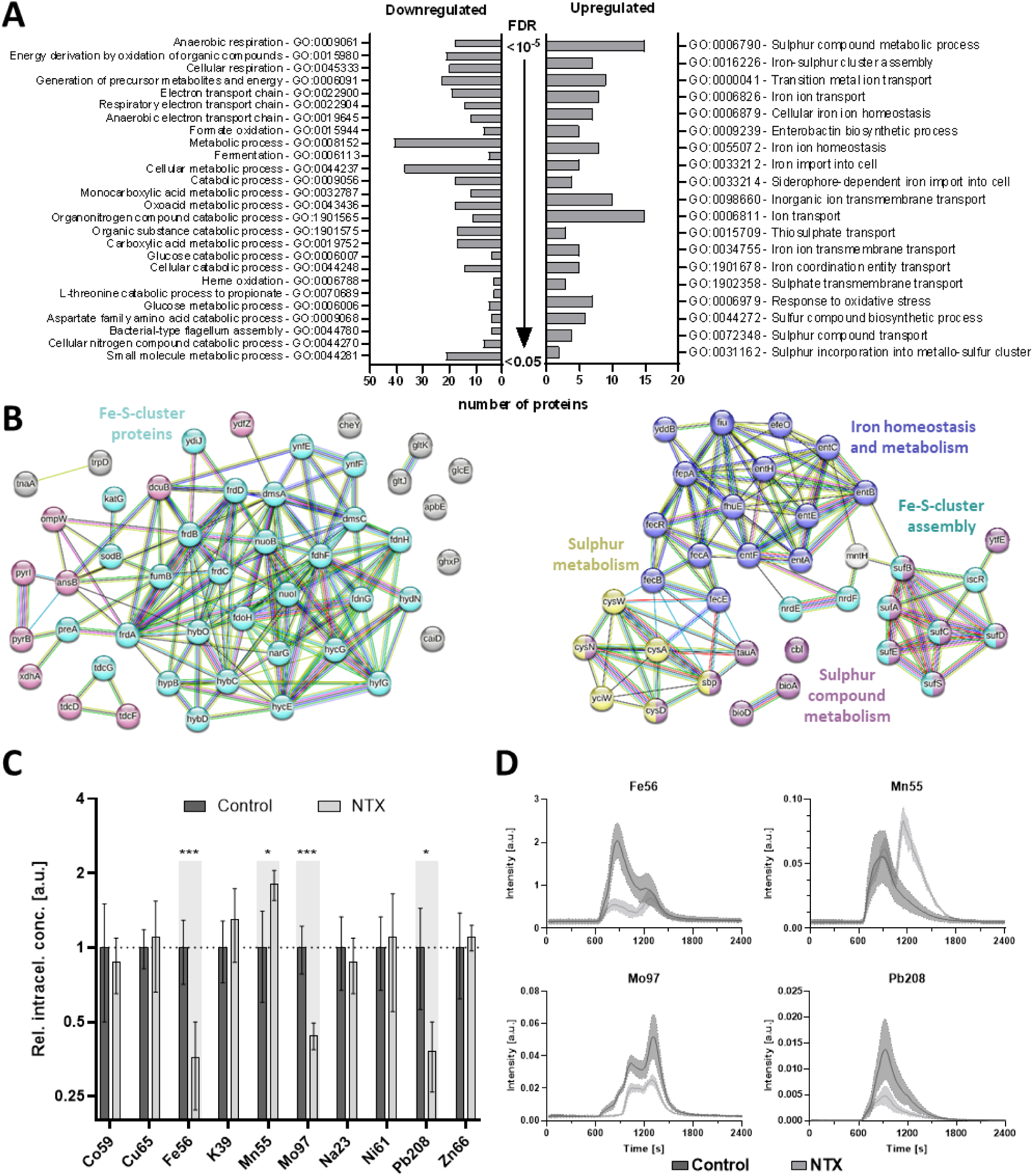
NTX treatment induces iron-deficiency in E. coli ATCC25922. A) Gene Ontology term (biological function) analyses of NTX treated E. coli with downregulated proteins (log2 < -2.5) to the left and upregulated proteins (log2 > 1.5) to the right. The number of proteins assigned to the respective GO term is displayed, with GO-terms sorted from lowest false discovery rate (FDR) on the top to highest FDR (while still being significant) at the bottom. B) String networks of respective down- (left) or upregulated (right) proteins coloured according to their assigned group. For downregulated proteins, light-blue proteins require Fe-S-clusters for function, while pink proteins show a direct connection to Fe-S-cluster proteins. Upregulated proteins are assigned to iron homeostasis (blue), sulphur (compound) metabolism (yellow and/or violet, respectively) and Fe-S-cluster assembly (light-blue). Proteins not assigned to any group are grey. C) Relative intracellular elemental concentration (SEC-ICP-MS) reveals lack of protein-bound iron, molybdenum and lead in NTX-treated E. coli (light grey), while manganese concentration increased. For quantification, peak areas were summed up, normalized against DMSO-treated control cells (dark grey), and compared using t-tests (n = 4, mean and SD are shown). D) Individual SEC-ICP-MS chromatograms of significantly affected elements. The thick line represents the mean of four independent samples, while the dashed line (filled area) represents the SD. For smoothing, the moving average of five neighbouring data points was displayed.

The overall cellular response to NTX-induced stress was characterised by a drastic downregulation of respiration- and metabolism-related proteins, while all significantly upregulated proteins could be assigned to iron homeostasis, sulphur metabolism and Fe-S-cluster assembly (**Figure 2**A/B). Among them are those involved in siderophore transport and synthesis (Enterobactin, 4.2-fold), Fe-citrate transport (Fec system, 4.4-fold) and proteins expressed during iron starvation like NrdF (32.5-fold) and NrdE (11-fold) (Martin & Imlay, 2011). Over 60% of all significantly less abundant proteins (29/47) require Fe-S-cluster proteins for function, while more than 80% (38/47) were directly connected to Fe-S-cluster proteins, according to String Network (**Figure 2**B). Both the fumarate reductase complex (FrdABCD, 15-fold) and the nitrate reductase (NarG, 7.5-fold) were strongly downregulated. A subsequent KEGG-pathway analysis of the same protein sets using Voronoi plots (Liebermeister *et al*, 2014) clearly confirmed a strong overall impact on cellular metabolism, as the majority of up- or downregulated proteins could be assigned (**Figure SI 11**).

SEC-ICP-MS enabled the characterization of protein-associated metal species in *E. coli*. In this setup, larger protein complexes eluted earlier (700–900 s), followed by smaller proteins (around 1,200 s), and finally low-molecular weight or unbound metal ions (> 3,000 s). NTX treatment led to significant changes in the elemental composition of *E. coli* (**Figure 2**C/D), **Figure SI 12**). For iron (Fe56), a marked reduction in the main protein-associated peak eluting at 800–900 s was observed, corresponding to a relative intensity of only 0.36 compared to control samples. Similar decreases were detected for Mo97 and Pb208, with relative intensities reduced to 0.44 and 0.38, respectively. In contrast, NTX exposure induced the appearance of a new manganese (Mn55) peak at approximately 1,200 s, resulting in a 1.82-fold increase in relative concentration compared to untreated cells. No significant changes were detected for cobalt (Co59), copper (Cu65), zinc (Zn66), or other elements (**Figure SI 12**).

The proteomic signature of NTX-treated *E. coli* clearly showed significantly affected cellular respiration, impaired bacterial energy production and ultimately a response to iron-deficiency on the biological level in *E. coli* (McHugh *et al*, 2003). Hence, we hypothesised that NTX’s antibacterial activity eventually causes iron deprivation in *E. coli*, and we, therefore, assessed the activity of NTX against *E. coli* strains in iron-limited medium (IL-MHBII).

We tested the KEIO library strain *E. coli* K12 Δ*entA*, deficient in enterobactin synthesis, and its respective wildtype strain *E. coli* K12/BW25113 in quadruplicates (**Table 1**). In regular MHBII medium, NTX MIC against both strains was found to be 8 µg/mL, whereas NTX activity in IL-MHBII was increased 2-fold against wildtype (4 µg/mL), and 2- to 4-fold against the *entA* knockout (2-4 µg/mL).

**Table 1:** Iron-deficiency renders E. coli more susceptible to NTX. Strains were tested in quadruplicates (A-D).

| Strain | Medium | MIC [$\mu\text{g/mL}$] | | | | |
| --- | --- | --- | --- | --- | --- | --- |
|  |  | A | B | C | D | Range |
| *E. coli* K12 BW25113 | MHBII | 8 | 8 | 8 | 8 | 8 |
|  | IL-MHBII | 4 | 4 | 4 | 4 | 4 |
|  | IL-MHBII + 1 mM Fe(III)-EDTA | 8 | 8 | 8 | 8 | 8 |
| | IL-MHBII + 700 $\mu\text{M}$ Zn<sup>2+</sup> | 32 | 32 | - | - | 32 |
| *E. coli* K12 $\Delta\text{entA}$ | MHBII | 8 | 8 | 8 | 8 | 8 |
|  | IL-MHBII | 2 | 2 | 4 | 2 | 2-4 |
|  | IL-MHBII + 1 mM Fe(III)-EDTA | 8 | 8 | 8 | 8 | 8 |

IL-MHBII: iron-limited medium following treatment with Chelex-resin

Subsequent re-supplementation of IL-MHBII with 1 mM Fe(III)-EDTA voided these effects and the MIC returned to baseline (8 µg/mL) for both strains confirming an influence of iron on NTX MoA. Interestingly, when we added 700 µM ZnCl_2_ to the testing medium, NTX activity was diminished 8-fold compared to no Zn^2+^-supplementation, suggesting that NTX saturation by *e.g*., zinc reduces its antibacterial effect. **NTX-metabolites are biologically inactive.** In humans, NTX is rapidly metabolised in the liver to its major metabolites, NTX-S and NTX-G (Wagenlehner *et al*, 2014) (**Figure 3**A), yet the contribution of these metabolites to NTX’s pharmacological effects remained unclear. To address this question, we, for the first time, synthesised these metabolites and were thus able to systematically analyse their activity. To generate the NTX-metabolites, NTX was sulphated with sulphur trioxide triethylamine complex (SO₃·NEt₃) to yield NTX-S as white amorphous solid after purification via C18 reversed-phase column chromatography (**Scheme SI 1**). NTX-G was obtained in a four-step synthesis from D-glucurono-3,6-lactone (**Scheme SI 2**). NMR and LC-MS analysis confirmed the absence of free NTX, consistent with the colourless appearance of NTX-G, in contrast to the yellow colour of free NTX.

**Figure 3:**
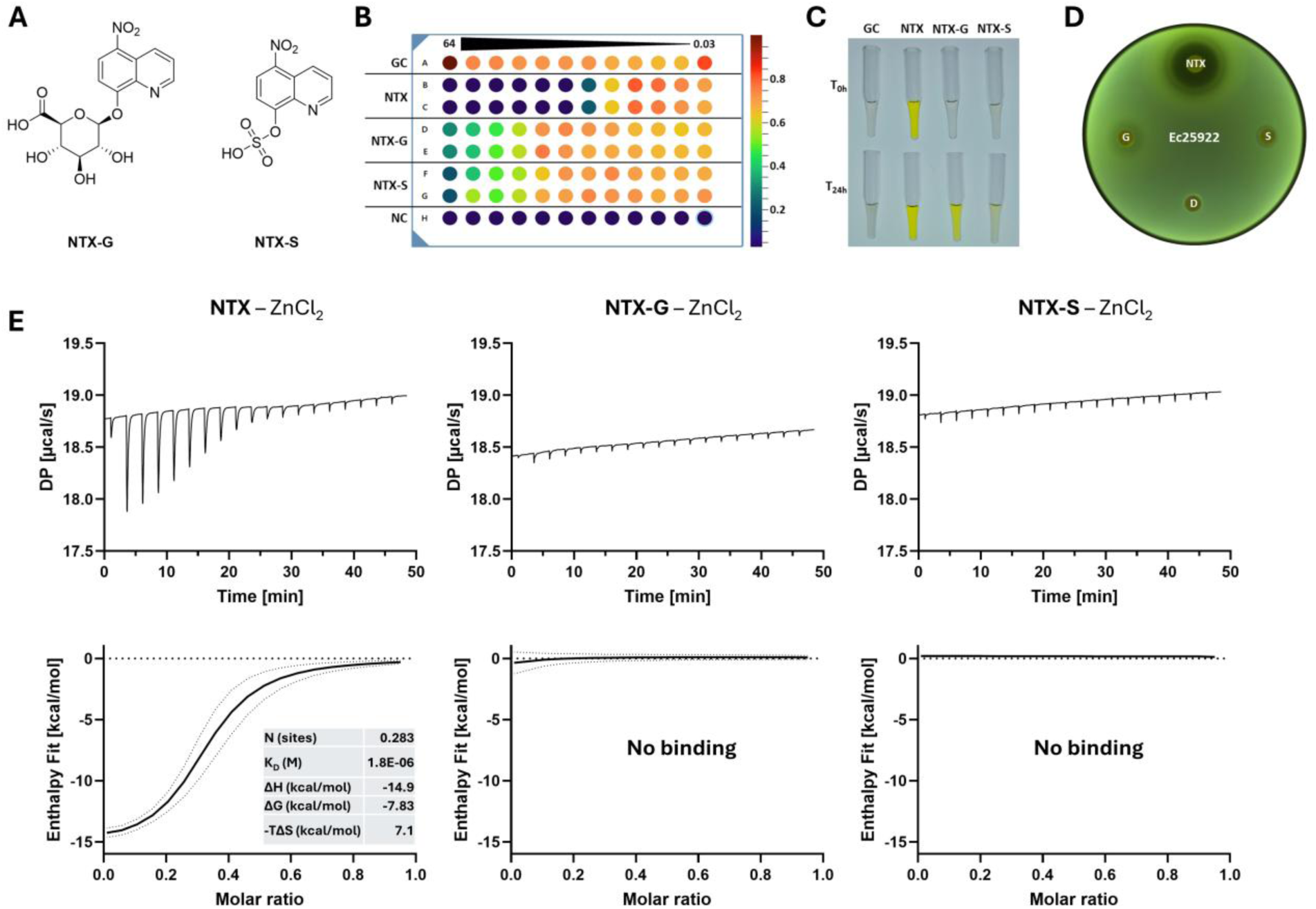
NTX metabolites NTX-S and NTX-G do not have antibacterial or chelating properties. A) Chemical structures of NTX metabolites. B) Standard broth micro dilution of NTX and its metabolites against E. coli ATCC25922; compounds are added at 64 µg/mL into the first column and sequentially diluted 1:2 until column twelve. The first row served as growth control (GC), while the last row contained only medium (negative control, NC). After incubation, OD600 was read and is displayed as colours from dark blue (no growth) to dark red (strong growth). C) 64 µg/mL of NTX and its metabolites incubated for 24 h with or without E. coli ATCC25922 shows increase in yellow colour in metabolite samples where bacteria were added. D) Disc diffusion with NTX, NTX-G (G), NTX-S (S), and DMSO (D), against E. coli ATCC25922. E) Isothermal titration calorimetry of NTX (n = 3) and metabolites (n = 2) against ZnCl2 reveals no binding interaction of metabolites with Zn^2+^.

Having both metabolites in hand, we performed standard susceptibility assays using *E. coli* ATCC25922 and also analysed the human liver cell line HepG2 for potential cytotoxic effects (**Figure 3**B-D,

**Table SI *2***). Although our primary focus was to better understand the antibacterial effects of NTX, we evaluated the cytotoxicity of its main metabolites, NTX-S and NTX-G, with NTX included as a reference compound, since the basis for its broad in vitro activity and its apparent lack of in vivo safety concerns remain poorly understood. While NTX resulted in the expected minimal inhibitory concentration (MIC) of 4 µg/mL and showed an IC_50_ of 1 µg/mL for E. coli and HepG2, respectively, both metabolites were found inactive (MIC > 64 µg/mL, IC_50_ HepG2 > 64 µg/mL). Bacterial cell growth could be observed even in wells containing the highest concentration of metabolite, although slight growth inhibition of E. coli was observed at these concentrations alongside a yellow colour. The observed colour change was stronger for NTX-G compared to NTX-S (**Figure 3**C). In a disc diffusion assay (**Figure 3**D), a clear zone of inhibition could be seen around the disc containing NTX, while no clear zones could be observed for the metabolites, confirming the lack of antibacterial activity of NTX metabolites. It has to be noted that again a slight halo was visible around metabolite discs with a mild yellow colouration that was not visible at the beginning of the experiment (for interpretation of this result see below).

Given that the molecule’s activity is presumed to arise solely from its metal-chelating properties, we conducted bacteria-free ITC experiments using Zn^2+^ and NTX metabolites. Zn²⁺ was used as a surrogate metal ion in ITC experiments due to its greater chemical stability and experimental reproducibility relative to Fe²^+^. As expected, neither NTX-S nor NTX-G exhibited direct interaction with Zn^2+^, confirming the absence of complex formation. In the same set up, a clear complex formation could be observed for NTX ([NTX_2_M]). (**Figure 3**E).

Thus, for the first time, we could confirm that NTX metabolites are biologically inactive, with no or only marginal effects on HepG2 and *E. coli* cell growth, and do not possess metal-chelating properties, raising the question why NTX during treatment of uUTI shows activity in the human bladder as current literature states that only 1% of NTX parent drug is found in urine, whereas NTX-G accounts for 55.7% and NTX-S for 38.4% (Wagenlehner *et al*, 2014).

### NTX metabolites are converted to active NTX in presence of various urinary pathogens

As the metabolites were inactive and colourless, we assessed whether biotransformation into the active compound NTX occurred, since the yellow coloration observed in our susceptibility assays strongly indicated NTX presence. To this end, 1 µM of each metabolite was incubated at 37 °C for 24 hours in pooled human urine (*ex vivo*) with or without *E. coli* ATCC25922 (OD_600_ 0.5); samples were taken at indicated time points (*n* = 3), followed by MS analysis of parent metabolite and resulting NTX (**Figure 4**A).

**Figure 4:**
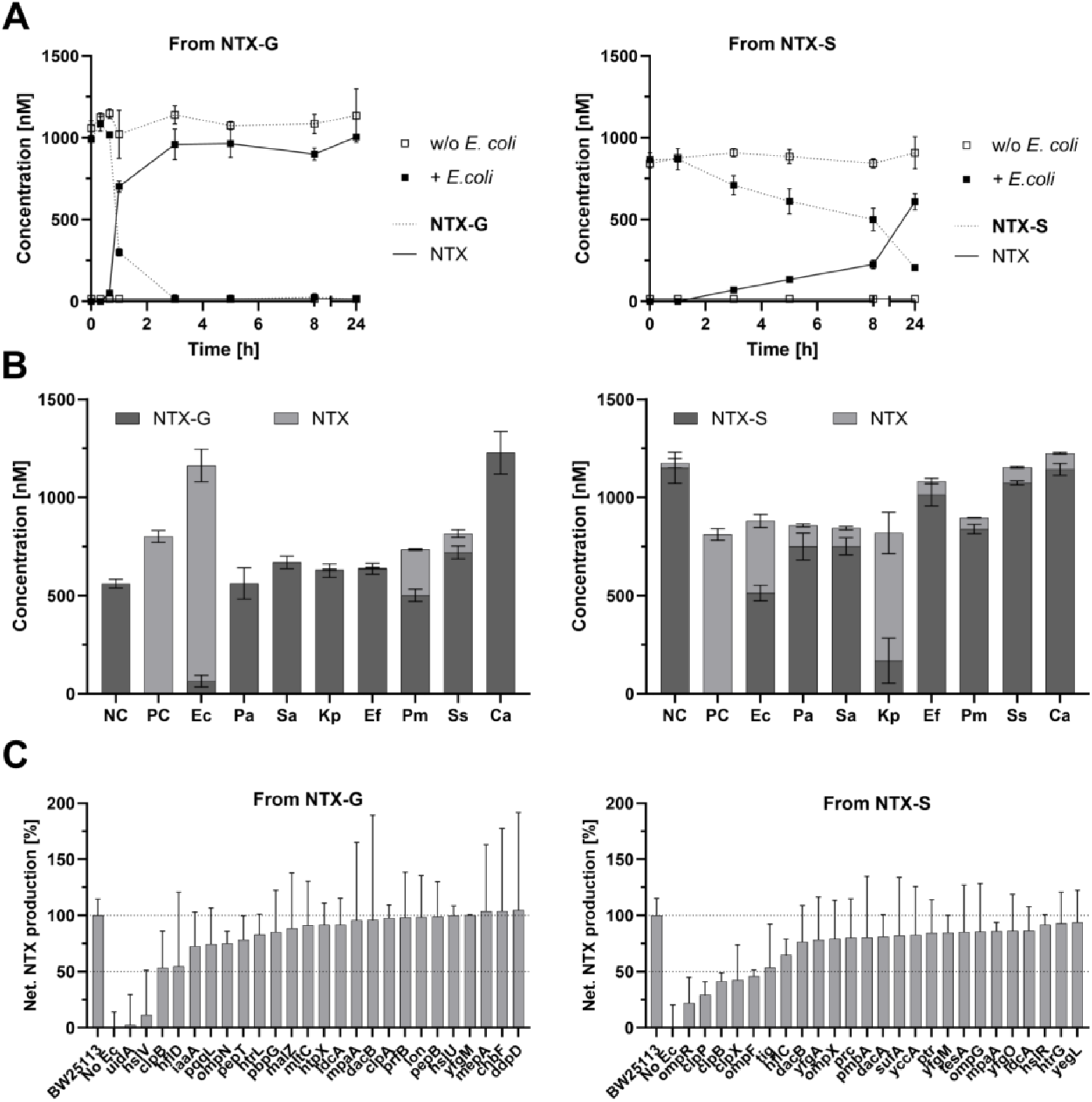
NTX metabolites can be converted to NTX in presence of bacteria. A) 1 µM NTX-G (left) or NTX-S (right) was incubated at 37 °C in presence (filled square) or absence (open square) of E. coli ATCC25922 (OD600 0.5) in human urine. At indicated time points, samples were taken and extracted for quantitative analysis of parent metabolite (dashed line) and NTX (solid line), mean and SD are shown (n = 3). B) NTX metabolite conversion by fresh UTI isolates in urine. Experiment was done twice with mean and SEM displayed. Medium w/o bacteria was used as negative control (NC), while 1 µM NTX was displayed as intensity reference (PC). Ec: E. coli, Pa: P. aeruginosa, Sa: S. aureus, Ss: S. saprophyticus, Kp: K. pneumoniae, Ef: E. faecalis, Pm: P. mirabilis, Ca: C. albicans. C) KEIO library screen reveals E. coli knockout strains unable to convert NTX metabolites. Selection of top 25 strains with lowest NTX production level are shown, refer to Figure SI 15 for all strains. NTX peak area was normalised against peak area of library wildtype E. coli BW25113, while NTX area in control samples (w/o bacteria) was set to zero.

Incubation of *E. coli* ATCC25922 with NTX-G and NTX-S demonstrated that the bacteria efficiently converted both metabolites into the active compound NTX, while no conversion was detected in control samples lacking bacterial cells. NTX-G was rapidly transformed, with only 30% NTX-G remaining after 1 h, and complete conversion after 3h. At the same time, NTX concentration increased concomitantly. In contrast, NTX-S was converted more slowly, reaching approximately 25% conversion after 8 h and about 60% after 24 h.

To assess species specificity and clinical relevance of this finding, we expanded the test panel to include freshly isolated clinical strains (21x *E. coli*, 8x *Pseudomonas aeruginosa*, 4x *Staphylococcus aureus*, 2x *Staphylococcus saprophyticus*, 7x *K. pneumoniae*, 6x *Enterococcus faecalis*, 7x *Proteus mirabilis*, 8x *Candida albicans*), obtained from UTIs in hospitalised patients. All isolates were incubated individually in urine supplemented with 1 µM metabolite and after 24 h, the presence of NTX was quantified. The species summary can be seen in **Figure 4**B and individual isolates in **Figure SI 13** (NTX-G) and **Figure SI 14** (NTX-S).

On average, *E. coli* isolates converted 91.6% of NTX-G to NTX, followed by 31.7% conversion by *P. mirabilis* and 11.8% by *S. saprophyticus*. Notably, 16 of the 21 *E. coli* isolates achieved > 95% NTX-G conversion, while only one isolate (Ec4) did not produce NTX from NTX-G at all. In contrast, NTX-S was metabolised by all tested species, with the highest activities observed in *K. pneumoniae* (79.4%) and *E. coli* (41.8%). The remaining species exhibited only low conversion rates of 7-12%, including one *Klebsiella* isolate (Kp1) that did not convert NTX-S at all.

To gain insights into potential genetic factors driving metabolite conversion, we selected over 100 strains from an *E. coli* transposon library (KEIO) carrying knockouts in various catabolic enzymes and exposed them to NTX metabolites. After 24 h incubation in urine (*n* = 2), we measured the presence of NTX in each sample and compared against the production of library wildtype *E. coli* BW25113 set to 100% and a urine control without bacteria used for background subtraction; the 25 strains with the lowest amount of NTX produced are displayed in **Figure 4**C, while all strains can be found in **Figure SI 15**. For NTX-S, all tested knockout strains retained some capacity to produce NTX, although strains lacking OmpR (22%), ClpP (29%), ClpB (42%), ClpX (43%), and OmpF (46%) showed markedly reduced conversion levels below 50% of wildtype NTX levels. In contrast, loss of UidA nearly abolished NTX-G conversion (2.6%), consistent with presumed direct hydrolysis of the glucuronide moiety by the β-glucuronidase. Loss of HslV also strongly impaired NTX-G conversion (11%), although whether this reflects a direct catalytic role or an indirect effect on cellular physiology remains elusive. The identification of UidA and HslV is notable because both proteins are localized in the bacterial cytoplasm. Their involvement therefore implies that NTX-G gains access to the intracellular compartment prior to conversion. While the uptake mechanism remains unknown, passive diffusion, porin-mediated transport, or uptake through carbohydrate transport systems may contribute and warrant further investigation. To validate the gene deletion of the KEIO library strain, absence of *uidA* was confirmed using direct colony PCR with gene specific primers (**Figure SI 16**). Subsequently, all *E. coli* isolates were analysed using the same set up revealing that all isolates but Ec4 contain at least one copy of *uidA* in their genome, perfectly matching the conversion data (**Figure SI 17**). To check for the presence of these genes across thousands of *E. coli g*enomes, we retrieved all available GenBank-annotated *E. coli* genomes from NCBI (*n* = 5,089) and used them as blast reference database for the *uidA* and *hslV* sequences. We confirmed that both genes belong to the core genome of *E. coli* and are present (sequence coverage > 95%) in 98% (*uidA*) and 100% (*hslV*) of all tested strains (**Figure SI 18**).

## Discussion

In this study, we applied a systems-biology approach to analyse the antibacterial activity of nitroxoline (NTX), revealing new insights into its metabolic fate and MoA with important implications for its clinical use. Across proteomic, elemental, and phenotypic assays, NTX treatment of *E. coli* reproducibly resulted in a response pattern consistent with iron starvation, likely affecting energy metabolism and cellular respiration. In addition, we show that the major urinary metabolites NTX-S and NTX-G lack intrinsic antibacterial activity. After their first chemical synthesis, we found them to be biologically inactive, but demonstrate that they can be reactivated via pathogen-dependent biotransformation in human urine, regenerating active NTX.

Proteome profiling revealed a reduced abundance of respiration-associated enzymes, including the fumarate reductase and nitrate reductase complexes, alongside a strong depletion of Fe–S-cluster-dependent proteins. In parallel, proteins involved in iron uptake, sulphur metabolism, and Fe–S-cluster assembly were among the ones significantly increased in abundance. This constellation of protein expression changes is a well-established bacterial response to impaired iron availability, irrespective of the upstream cause (Cacace *et al*, 2025; McHugh *et al*, 2003). The ribonucleotide reductase NrdEF (log_2_ = 3.48 and 5.02), which enables cell replication during iron starvation (Martin & Imlay, 2011), was over 30-fold upregulated in our experiments. Consistently, SEC-ICP-MS analysis demonstrated a pronounced loss of protein-bound iron in NTX-treated cells, while whole-cell protein levels remained comparable, indicating a specific perturbation of metal cofactor availability rather than general cytotoxicity. Iron-sulphur cluster and iron-homeostasis dysregulation have been described for copper- and zinc-induced metal toxicity, potentially suggesting that NTX acts as a metallophore resulting in a toxic intracellular accumulation of metal ions (Cacace *et al*, 2025; Chillappagari *et al*, 2010; Xu *et al*, 2019). In previous studies, we have reported EnvZ, OmpF, OmpC and LamB dysregulation in NTX-resistant *E. coli* mutants indicative for high cellular osmolarity (Deschner *et al*, 2024), which supports such theory and is also in line with recent insights into NTX’s MoA (Cacace *et al*, 2025). We further observed an increase in protein-bound manganese and a lack of molybdenum, which also supports phenotypic iron-deprivation. Manganese was shown to be important especially as a substitute for iron during oxidative stress or to compensate when iron is scarce, while molybdenum cofactor biosynthesis relies on iron, as key enzymes in this pathway require iron-sulphur ([4Fe-4S]) clusters for function (Čapek & Večerek, 2023; Zupok *et al*, 2019).

At the phenotypic level, iron limitation rendered *E. coli* more susceptible to NTX, while iron re-supplementation restored baseline susceptibility. Importantly, the data demonstrate that NTX-treated cells experience a certain functional iron deficit that directly compromises cellular respiration and energy generation, which is ultimately incompatible with sustained bacterial growth, as energy generation becomes inefficient (Baez *et al*, 2022). The underlying molecular origin of this iron deprivation cannot be conclusively resolved based on the present data. NTX is a potent chelator of several transition metals, including Fe²⁺, Cu²⁺, and Zn²⁺, as confirmed here by quantitative ITC analysis. Direct sequestration of iron by NTX might therefore also be a plausible contributing mechanism. At the same time, NTX has been described to act as a metallophore for copper and zinc in other bacterial species, leading to intracellular metal accumulation and subsequent destabilisation of Fe–S-cluster proteins (Cacace *et al*, 2025). Such metal-induced Fe–S-cluster instability is also known to trigger iron-starvation responses at the transcriptional and post-transcriptional level (Chillappagari *et al*, 2010; Xu *et al*, 2019).

Our data do not allow us to distinguish whether the observed iron deprivation arises predominantly from direct iron chelation, from metal-induced Fe–S-cluster damage, or from a combination of both processes. Regardless of the proximal cause, the convergent consequence is sequestration of functional iron from essential respiratory and metabolic enzymes. The bacterial cell attempts to compensate by upregulating iron acquisition and alternative ribonucleotide reductases (Martin & Imlay, 2011), but this response is insufficient to restore energy homeostasis. We therefore conclude that iron deprivation represents the unifying downstream mechanism that explains NTX antibacterial activity in *E. coli*.

This interpretation also reconciles our findings with previous reports describing metal-dependent modulation of NTX activity (Sobke *et al*, 2012). Saturation of NTX coordination sites with excess zinc abolished antibacterial activity, consistent with metal chelation being essential for NTX function. By contrast, magnesium and calcium did not bind NTX directly in our assays and did not show a consistent impact on activity, indicating that previously reported effects of NTX on RNA polymerase are likely indirect (Pelletier *et al*, 1995; Fraser & Creanor, 1974). Furthermore, our proteomic signature does not resemble that of classical RNA polymerase inhibition (Meneguello *et al*, 2020). Overall, our data support a model in which NTX exerts its antibacterial effect through metal coordination, with iron deprivation as the decisive physiological outcome leading to respiratory collapse and growth inhibition.

The iron-deprivation signature observed for NTX-treated *E. coli* is consistent with a metal-chelation-based mechanism of antibacterial activity. However, this is difficult to reconcile with human pharmacokinetics, as NTX is rapidly metabolised and renally excreted predominantly as sulphate and glucuronide conjugates (NTX-S and NTX-G) (Wagenlehner *et al*, 2014; Wijma *et al*, 2018; Forstner *et al*, 2018). Because these modifications block the 8-hydroxy group required for metal coordination, a key question is how NTX retains clinical efficacy despite extensive metabolic inactivation. We demonstrate for the first time that the major NTX metabolites NTX-S and NTX-G are biologically inactive. Both conjugates lacked antibacterial and cytotoxic activity and failed to bind divalent metal ions as exemplified by Zn^2+^ binding experiments. Modification of the 8-hydroxy group abolishes the essential electron-donating site required for metal coordination, thereby eliminating chelator activity. It has already been reported for glucoconjugates of NTX (8-quinolinyl-β-D-glucopyranoside) and clioquinol (5-chloro-7-iodo-8-quinolinyl-β-D-glucopyranoside), synthesised as potential anti-cancer compounds, that glucose addition at this hydroxyl group significantly diminishes antiproliferative activity against cancer cell lines leading the authors to conclude that the presence of a sugar moiety in a prodrug approach could eliminate the undesired chelation of systemic metal ions (Oliveri *et al*, 2012). Our findings demonstrate that NTX metabolites cannot contribute to antibacterial activity in their conjugated form. This result creates an apparent paradox: Clinically, NTX is effective in urinary tract infections, yet the active parent compound is largely absent from urine. Our data resolve this paradox by demonstrating pathogen-driven reactivation of NTX metabolites at the site of infection. In *ex vivo* urine assays, clinically relevant uropathogens, most prominently *E. coli* and *K. pneumoniae*, efficiently reconverted NTX-G and, to a lesser extent, NTX-S into active NTX. Importantly, this transformation did not occur in sterile urine, demonstrating that it requires the presence of viable uropathogenic bacteria.

Genetic analysis in *E. coli* identified the β-glucuronidase UidA as the principal determinant of NTX-G conversion, establishing a direct enzymatic basis for metabolite reactivation consistent with previous reports on glucuronide hydrolysis of compounds such as arbutin and 8-hydroxyquinoline (Oliveri *et al*, 2012; Parejo *et al*, 2001). The near-complete loss of conversion in the *uidA* mutant strongly supports a direct role of β-glucuronidase-mediated hydrolysis of NTX, whereas the contribution of the protease subunit HslV is less clear. The identification of UidA also raises an important mechanistic question regarding metabolite uptake. As UidA is a cytoplasmic enzyme, NTX-G must access the intracellular compartment prior to hydrolysis, while the transport pathway remains unknown. In contrast, NTX-S conversion appeared less specific and was only partially reduced in mutant strains lacking components of the Clp protease system or the OmpF/OmpR porin regulatory network, suggesting a more complex or indirect mechanism. The Clp system is essential for proteolytic activity of bacterial cells, whereas OmpF and OmpR contribute to outer membrane permeability and may therefore influence metabolite uptake (Gerken *et al*, 2020; Aljghami *et al*, 2022). Notably, we previously identified *envZ* mutations in NTX-resistant *E. coli* resulting in altered OmpF/OmpR expression in these mutants (Deschner *et al*, 2024). The reduced conversion of NTX-S observed in *ompF* and *ompR* mutants is therefore consistent with a model in which membrane permeability contributes to metabolite uptake prior to intracellular deconjugation. Whether NTX metabolites enter the cell via porin-mediated diffusion, specific transporters, or alternative uptake pathways remains an important subject for future investigation. The inability of one *Klebsiella* isolate to convert NTX metabolites further suggests that additional, potentially more critical, determinants of metabolite reactivation remain to be identified. A broader analysis of clinical isolates may reveal genetic determinants of metabolite reactivation and establish whether bacterial conversion capacity can be leveraged as a biomarker of NTX treatment efficacy.

To our knowledge, this is the first experimental demonstration that NTX undergoes enzymatic reactivation from its Phase II metabolites by bacterial pathogens at the site of infection. While analogous prodrug-like activation mechanisms have been described for other glucuronide conjugates (Oliveri *et al*, 2012; Parejo *et al*, 2001), this concept had not previously been experimentally addressed for NTX. Importantly, in the present case, metabolite formation is not a drug design feature but the result of host detoxification. Bacterial reactivation effectively inverts this process, selectively restoring antibacterial activity only where pathogens are present. This host–pathogen interplay provides a coherent explanation for several defining features of NTX pharmacology. Systemic detoxification through rapid conjugation limits host toxicity despite the potent metal-chelating properties of the parent compound. At the same time, pathogen-specific reactivation ensures local appearance of active NTX in infected urine, where metal coordination leads to iron deprivation and bacterial growth inhibition. In this framework, NTX metabolites function as inactive precursors rather than as direct effectors, and bacterial metabolism becomes a prerequisite for therapeutic efficacy.

The requirement for bacterial biotransformation may explain the low levels of free NTX in urine samples that were taken from healthy individuals (Wijma *et al*, 2018), but further pharmacokinetic studies are necessary to improve our understanding of the conversion rate in humans, especially those suffering from an active UTI. Our proof-of-concept experiments *ex vivo* show almost quantitative conversion of NTX metabolites in the presence of ∼10⁸ CFU/mL *E. coli* within a few hours, a concentration exceeding typical bacterial loads during UTIs (>10⁵ CFU/mL) (Hay *et al*, 2016). Despite the obvious requirement for metabolic activation, NTX has demonstrated robust antibacterial activity across decades of use, underscoring its real-world efficacy, hence, additional potential contributing factors such as the urobiome should also be considered (Brubaker *et al*, 2021).

The activation-at-site mechanism described here may also contribute to the favourable clinical profile of NTX. Recent studies support its efficacy and tolerability in long-term prophylaxis of recurrent UTIs (Wiedemann *et al*, 2026). Usually, antibiotic prophylaxis is associated with resistance concerns and contradicts the recommendations to reduce the usage of antibiotics as much as possible (Dhole *et al*, 2023; Global Antimicrobial Resistance and Use Surveillance System (GLASS) Report 2022, 2022). However, NTX possesses several characteristics that may favour prophylactic use, including low resistance rates, reduced fitness of resistant mutants, favourable tolerability, and, as demonstrated here, systemic conversion into inactive metabolites that require bacterial reactivation for antibacterial activity (Deschner *et al*, 2024; Stoltidis-Claus *et al*, 2023; Wiedemann *et al*, 2026; Naber *et al*, 2014). Ultimately, with our findings we potentially open the possibility for entire new applications for NTX that certainly underlines its clinical potential, even decades after its market approval.

In summary, our data support a model in which NTX antibacterial activity arises from metal coordination leading to iron deprivation and respiratory failure, while clinical efficacy depends on pathogen-driven reactivation of inactive host metabolites at the site of infection. This activation-at-site principle represents a previously unrecognised component of NTX pharmacology and provides a mechanistic rationale for its long-standing efficacy, favourable safety profile, and low resistance emergence.

## Data Availability

Datasets and further information of current and ongoing related studies are available upon request from the corresponding author. The mass spectrometry proteomics data have been deposited to the ProteomeXchange Consortium via the PRIDE partner repository with the dataset identifier PXD060245.

## Materials and Methods

### Strains and cultivation

Strains were purchased from the American Type Culture Collection or from the KEIO library. They were propagated in regular cation-adjusted Müller-Hinton Broth (MHBII) or Lysogeny Broth (LB) at 37 °C, and plated on Caso agar. For KEIO strains, 50 µg/mL kanamycin was added for regular propagation. The clinical isolates stemmed from patients with urinary tract infections who underwent microbiological diagnostics of urine samples at Saarland University Medical Center in Homburg, Germany. The samples were randomly chosen from a diagnostic biobank. All samples were obtained as part of routine diagnostic care and no specific ethics approval was required for the intended analysis.

### Isothermal Titration Calorimetry

Experiments were performed using a NanoITC 2G (TA Instruments) equipped with a 1033 µL cell and a 250 µL syringe (TA Instruments, T140721) with its respective software ITCrun. Samples were prepared in fresh 100 mM Tris/HCl pH 6.2 (or as otherwise stated) supplemented with 10% DMSO. For regular set up, 800 µM NTX was filled in the cell (ligand), while 2.4 mM of respective cation (as chloride salt) was filled in the syringe (analyte); concentrations were adjusted when necessary/possible to obtain c values ranging from 10-1000, calculated using following formula: c = n·[M]_cell_/K_d_. Buffer titrations were performed to check for dilution heat. Stirring speed was set to 250 rpm, while the analyte was injected over 20-30 injections of 5 µL with adequate distance to allow for baseline separation. All experiments with detected binding were performed at least in triplicates at 25 °C. The cell and stirrer were thoroughly cleaned using both methanol and MQ water between each run. Data evaluation was done using the NanoAnalyze software (TA Instruments) using both the independent model and the blank (linear) model. For iron experiments, all solutions have been thoroughly purged using nitrogen-flow to reduce the amount of dissolved oxygen as much as possible to increase Fe^2+^ stability.

For metabolite binding studies, experiments were performed using a Malvern MicroCal PEAQ-ITC (Malvern Panalytical) equipped with a 200 µL cell and a 40 µL syringe with its respective software. Stirring speed was set to 750 rpm, while the analyte (500 µM) was injected over 1 injection with 0.4 µL, followed by 18 injections with 2 µL into NTX (100 µM). All metabolite experiments were performed in duplicates at 25 °C. The cell and stirrer were thoroughly cleaned using methanol, 10% Contrad 70 and MQ water between each run, using the automated cleaning module. Data evaluation was done using the MicroCal PEAQ-ITC software (Malvern Panalytical). Thermodynamic parameters were calculated from equation ΔG = ΔH − TΔS = RT ln K_A_ = −RT ln K_D_ where ΔG, ΔH, and ΔS are the changes in Gibbs free energy, enthalpy, and entropy of binding, respectively. T is the absolute temperature, and R = 1.98 cal mol^−1^ K^−1^.

### Full proteome analysis

Sample preparation was based on previously published methods (Kirsch *et al*, 2021). Per repeat, a liquid overnight culture of *Escherichia coli* ATCC25922 was inoculated (1:100) in 5 mL fresh media and treated with 0.25x MIC NTX, or the respective amount of DMSO as negative control. The liquid cultures were incubated at 37 °C, 180 rpm until they reached the early stationary growth stage. Subsequently, 1 mL sample was collected in an Eppendorf tube on ice, centrifuged (6000x *g*, 4 °C, 5 min) and washed with 1 mL cold PBS, before storing the pellet at -80 °C until further use. Collected samples were lysed by adding 210 µL 0.4% SDS in PBS and three rounds of sonication with an amplitude of 50% for 30 s (Bandelin Sonoplus). Subsequently, samples were centrifuged (16900x *g*, 20 min, RT) and the supernatant was used to determine the protein concentration using BCA assay (Thermo). Proteome concentrations of the samples were adjusted to 100 µg/200 µL per sample. Afterwards, the proteome was precipitated by adding 1 mL ice-cold acetone and incubation overnight at -20 °C. Next, the samples were centrifuged (16900xg, 15 min, 4 °C), and supernatant was discarded. Samples were washed two times with 1 mL ice-cold methanol. Preparation for protein digest was done by resuspension in 200 µL X-buffer (7 M urea, 2 M thiourea, 20 mM HEPES, pH = 7.5) followed by reduction of cysteine with 0.8 µL DTT (250 mM) (dithiothreitol) for 45 min at 25 °C, capping with 2 µL IAA (550 mM) (iodoacetic acid) for 30 min at 25 °C and adding additional 3.2 µL DTT (250 mM) for 30 min at 25 °C. The protein digest was performed by adding 600 µL ammonium bicarbonate buffer (pH 8.5) and trypsin (1 µg) (MS grade, Promega) and incubation overnight at 37 °C, 400 rpm shaking. Digestion was stopped by adding 8 µL formic acid. Peptide samples were desalted on SepPak C18 columns (Waters) using the following protocol. Columns were primed by adding 1 mL 100% and 80% acetonitrile (ACN) with 0.5% FA, followed by equilibration with 3x 1 mL H_2_O + 0.1% TFA, before samples were loaded and washed with 3x 1 mL 0.1% TFA and 1x 0.5 mL 0.5% FA. Subsequently, samples were eluted into 2 mL LoBind (Eppendorf) by adding 3x 250 µL 80% ACN + 0.5% FA under slight vacuum and dried using SpeedVac before solving in 1% FA with a proteome concentration of 1 µg/µL. Samples were filtered through 0.22 µm membrane (Merck Millipore, UFC30GVNB) and transferred to QuanRecovery autosampler vials (Waters). Sample analysis was done by using nanoElute nano flow liquid chromatography system (Bruker, Germany) coupled with a timsTOF Pro (Bruker, Germany). Samples were loaded to the trap column (Thermo Trap Cartridge 5 mm) and washed with 6 µL 0.1% FA with a flow rate of 10 µL/min. Peptides were then transferred to the analytical column (Aurora Ultimate CSI 25 cm x 75 µm ID, 1.6 µm FSC C18, IonOpticks) and separated by a gradient elution (eluent A: H_2_O + 0.1% FA, B: ACN + 0.1% FA; 0% to 3% in 1 min, 3% to 17% in 57 min, 17% to 25% in 21 min, 25% to 34% in 13 min, 34% to 85% in 1 min, 85% kept for 8 min) with a flow rate of 400 nL/min. Captive Spray nanoESI source (Bruker, Germany) was used to ionize the peptides at 1.5 kV with 180 °C dry temperature at 3 L/min gas flow. TimsTOF Pro (Bruker, Germany) was operated in default dia-PASEF long gradient mode with TIMS set to 1/K0 start at 0.6 Vs/cm2, end at 1.6 Vs/cm2 with a ramp and accumulation time of 100 ms each and a ramp rate of 9.43 Hz. Mass range was set from 100.0 Da to 1700 Da with positive ion polarity. Dia-PASEF mass range was set to 400.0 Da to 1201.0 Da with a mobility range of 0.60 1/K0 to 1.43 1/K0 and a cycle time of 1.80 s. Collision energy for 0.60 1/K0 was set to 20.00 eV and for 1.6 1/K0 to 59.00 eV. Tuning MIX ESI-TOF (Agilent) was used for calibration of *m/z* and mobility. Data were processed using DIA-NN (version 1.8.1), and proteins were identified against Uniprot *E. coli* K12 reference proteome (Proteome ID: UP000000625, downloaded 16/01/2023). Settings were used as default except precursor charge range was from 2 to 4. C carbamidomethylation was set as fixed modification. “--relaxed-prot-inf” was added in additional options to allow further data processing with Perseus Software. In Perseus (version 2.0.5.0) the values were transformed to their log_2_-value and the replicates were grouped. The results were filtered by valid values in 3 out of 4 in at least one group. Missing values were imputed by default settings and the differential protein abundances between different conditions were evaluated using two-tailed student’s t-test. Cut-off for –log_10_ p-value was set to 1.3 (p-value 0.05), and thresholds were set to [log_2_ > 1.5] and [log_2_ < -2.5] for proteins present in higher or lower abundance, respectively – for the ease of description, we refer to this as up- or downregulated proteins. Different thresholds were used as the number of downregulated proteins was very large, potentially resulting in inaccurate analyses and false positive clustering. For evaluation, GO terms and KEGG pathways were analysed using String Network (string-db.org) and Voronoi plots (proteomaps.net) (Liebermeister *et al*, 2014), together with protein information retrieved from UniProt (uniprot.org). **SEC-ICP-MS.** Sample preparation was performed as described for the full proteome analysis above. After cell harvesting, cell pellets were suspended in 10 mM Tris-HCl (pH 7.4) and proteins were extracted using bead beating (5 cycles of 30 seconds at 6.5 m/s^2^) using 0.1 mm diameter quartz beads. Insoluble material was removed by two consecutive centrifugation steps (5 min at 5,000x *g* and 10 min at 10,000x *g*, respectively; 4 °C both) and samples were stored at -20 °C until further use in single-use aliquots. Protein content in the supernatants was determined using Bradford assay against bovine serum albumin as external calibrant. To determine intracellular elemental concentrations, 20 µL sample were separated on a Superose 6 Increase 3.2x300 gel-filtration column by isocratic elution with 10 mM Tris-HCl at a flow rate of 100 µL/min. The eluate was directly infused into an Agilent 7500c ICP-MS instrument, equipped with a Scott type spray chamber and a PFA µ-flow nebuliser, to monitor the intensity for several elemental isotopes over a period of 120 min. The plasma was operated at 1600 W and all other parameters were optimised daily to obtain stable signals for the elements of a tuning solution (10 ppb of Li-6, Y-89, Ce-140, Tl-205), as well as stable background signals for C-13, Na-23, and K-39 when infusing the eluent from the gel-filtration column.

Using in-house R-scripts, the resulting chromatographic traces were normalised for the natural isotopic abundance and a heavily smoothed C-13 baseline to enable semi-quantitative comparison of elemental elution profiles. Peak detection and integration was performed in Fityk (v1.3.1) (Wojdyr, 2010) after scaling each chromatogram to the injected protein amount. For each condition, four biological replicates were analysed and differences in total elemental concentration were assessed by multiple unpaired Student’s t-tests based on the summed peak areas of the respective chromatograms.

### Synthesis of metabolites

For NTX-S, NTX was sulphated with sulphur trioxide triethylamine complex (SO₃·NEt₃) (**Scheme SI 1**), followed by quenching in HEPES solution, to give NTX-S as white amorphous solid after purification via C18 reversed-phase column chromatography. For the synthesis of NTX-G, the protected glucuronyl bromide 2 was obtained via NaOMe-mediated ring opening, followed by acylation and bromination (**Scheme SI 2**) (Herceg *et al*, 2018; Walther *et al*, 2019). Coupling with NTX under Koenigs-Knorr conditions afforded the protected conjugate 3 in moderate yield. A subsequent two-step deprotection (deacetylation, then ester saponification) minimised elimination side reactions to the α,β-unsaturated acid and afforded NTX-G. The products were analysed by NMR and LC-MS, confirming the absence of free NTX (yellow solid), which was consistent with the colourless appearance of NTX-G and NTX-S in both solid and solution state. Detailed procedures for the synthesis of compounds are provided in the Supplementary Information.

### Production of iron-limited medium

MHBII was used as basis for iron-limited medium (IL-MHBII) utilising Chelex Resin 100 (200-400 mesh, Biorad). For ion removal, 1 g of resin was added per 10 mL medium, and left on a shaker for 2 h. Subsequently, the resin was removed using a regular paper filter and the medium was re-supplemented with necessary cations (22.5 mg/L Ca^2+^, 11.25 mg/L Mg^2+^ and 1 mg/L Zn^2+^). The pH was adjusted to 7.4 prior sterilization via 0.2 µm filter.

### Susceptibility testing

Minimal inhibitory concentrations were determined using standard broth microdilution according to EUCAST guidelines (ISO20776-1:2019) in round-bottom 96-well plates, and evaluated using a Tecan Spark reader. Regular quality control was performed using *Escherichia coli* ATCC25922 and appropriate reference antibiotics. For disc diffusion assays, liquid agar was inoculated (OD_600_ 0.05) and 20 mL were used per plate. 30 µg of test substance dissolved in 6 µL DMSO (5 mg/mL) was spotted on sterilised filter discs prior placing on agar surface and incubation at 37 °C. Plates were documented using a colony counter (Interscience Scan 500).

### Biotransformation and mass spectrometry

Indicated strains were grown ON in LB medium, split 1/5 and grown to early logarithmic phase. Cultures were adjusted to OD_600_ 0.5 in pooled human urine (University Hospital Basel) supplemented with 1 µM metabolite and incubated at 37 °C (shaking). For high-throughput set ups (clinical isolates and KEIO Screen), samples were incubated in 96-well plates sealed with oxygen-seals (BreathEasy, Roth, T093) to avoid evaporation. At indicated time points, samples were drawn and diluted 1/3 into 15 nM diphenhydramine (DPH) in MeOH/ACN (10%) for extraction. Subsequently, samples were centrifuged and the supernatant was transferred to MS vials or stored at -80 °C until further analysis. For absolute quantification, standards were prepared in urine (3-fold concentration, 3 µM to 1.46 nM), and also extracted in DPH/MeOH/ACN, to reach an upper quantification limit of 1 µM.

The amount of NTX and its metabolites was analysed by HPLC-MS/MS (Vanquish Flex coupled to a TSQ Altis Plus, Thermo Fisher, Dreieich, Germany). HPLC conditions were as follows: column, Hypersil GOLD C18 (1.9 µm, 100 x 2.1 mm; Thermo Fisher), temperature, 40 °C, eluent A: H_2_O + 0.1% formic acid, eluent B: acetonitrile + 0.1% formic acid, flow: 0.7 mL/min. The gradient was set to 10% B from 0 to 0.2 min, 10-90% B from 0.2 to 0.5 min, 90% B from 0.5 to 1.6 min and 10% B from 1.6 to 2.0 min. MS parameters: Spray voltage 3000 V (positive mode) / 3280 (negative mode), vaporizer temperature 350 °C, ion transfer tube temperature 380 °C, sheath gas 30 (arb), auxiliary gas 10 (arb), ion sweep gas 2 (arb). Mass transitions for NTX, its metabolites and DPH are given in **Table 2**. Quantification was done using Xcalibur Quan Browser (Thermo Fisher) with DPH as internal standard to adjust for sample variation.

**Table 2:** NTX and metabolite quantification parameters for HPLC-MS/MS analysis.

| Compound | Polarity | Precursor<br>[*m/z*] | Product<br>[*m/z*] | Collision energy<br>[V] | RF lens<br>[V] | RT<br>[min] |
| --- | --- | --- | --- | --- | --- | --- |
| NTX | + | 190.845 | 126.917<br>144.917 | 36<br>20 | 57<br>57 | 1.24 |
| NTX-G | + | 366.967 | 144.883<br>190.967 | 42<br>16 | 55<br>55 | 1.10 |
| NTX-S | - | 268.883 | 158.967<br>188.967 | 29<br>15 | 44<br>44 | 1.14 |
| DPH | + | 256.050 | 164.967<br>166.967 | 42<br>13 | 30<br>30 | 1.18 |

### Colony PCR

Colonies were picked, resuspended in 0.9% saline solution, and microwaved for 5 min at 900 W to generate template DNA. A master mix was created using 12.5 µL One Taq Quick-Load 2X Master Mix (NewEngland Biolabs, M0486), 2 µL template, 0.5 µL of forward and revers primers (10 µM, UidA_F1: ATGTTACGTCCTGTAGAAACCCCAACCC; UidA_F2: GTAAGGGTAATGCGAGGTACGGTAGG), and 9.5 µL nuclease-free water (total of 25 µL per reaction). PCR cycler protocol was used as follows: 5 min 94 °C, 30 cycles of 30s/94 °C, 60s/60 °C, 60s/68 °C, followed by 5 min final extension at 68 °C and hold at 4 °C.

## Supplementary Information

**Figure SI 1:**
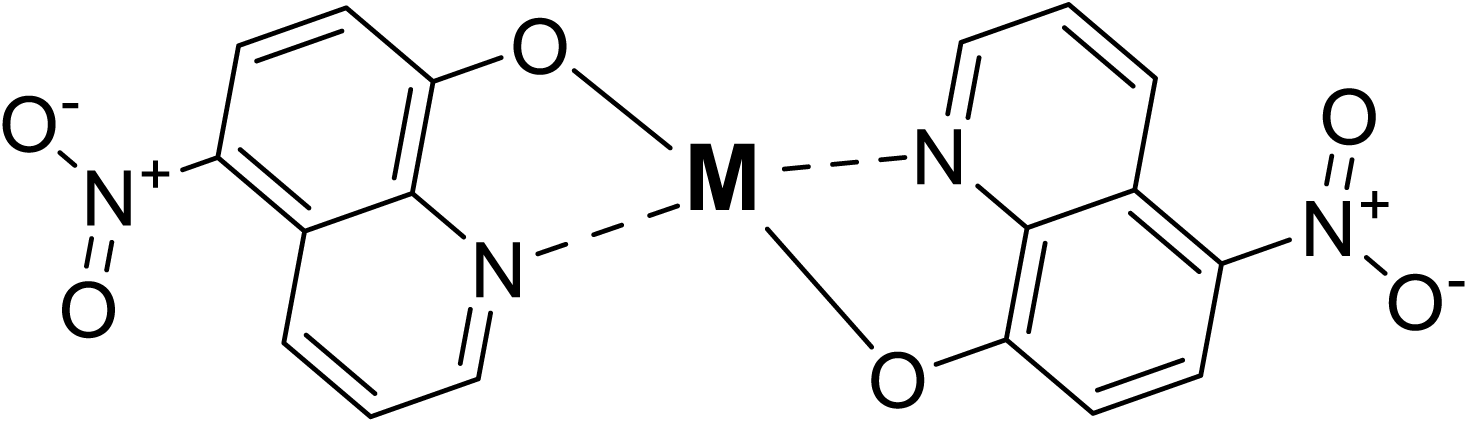
General structure of an NTX-metal complex.

**Figure SI 2:**
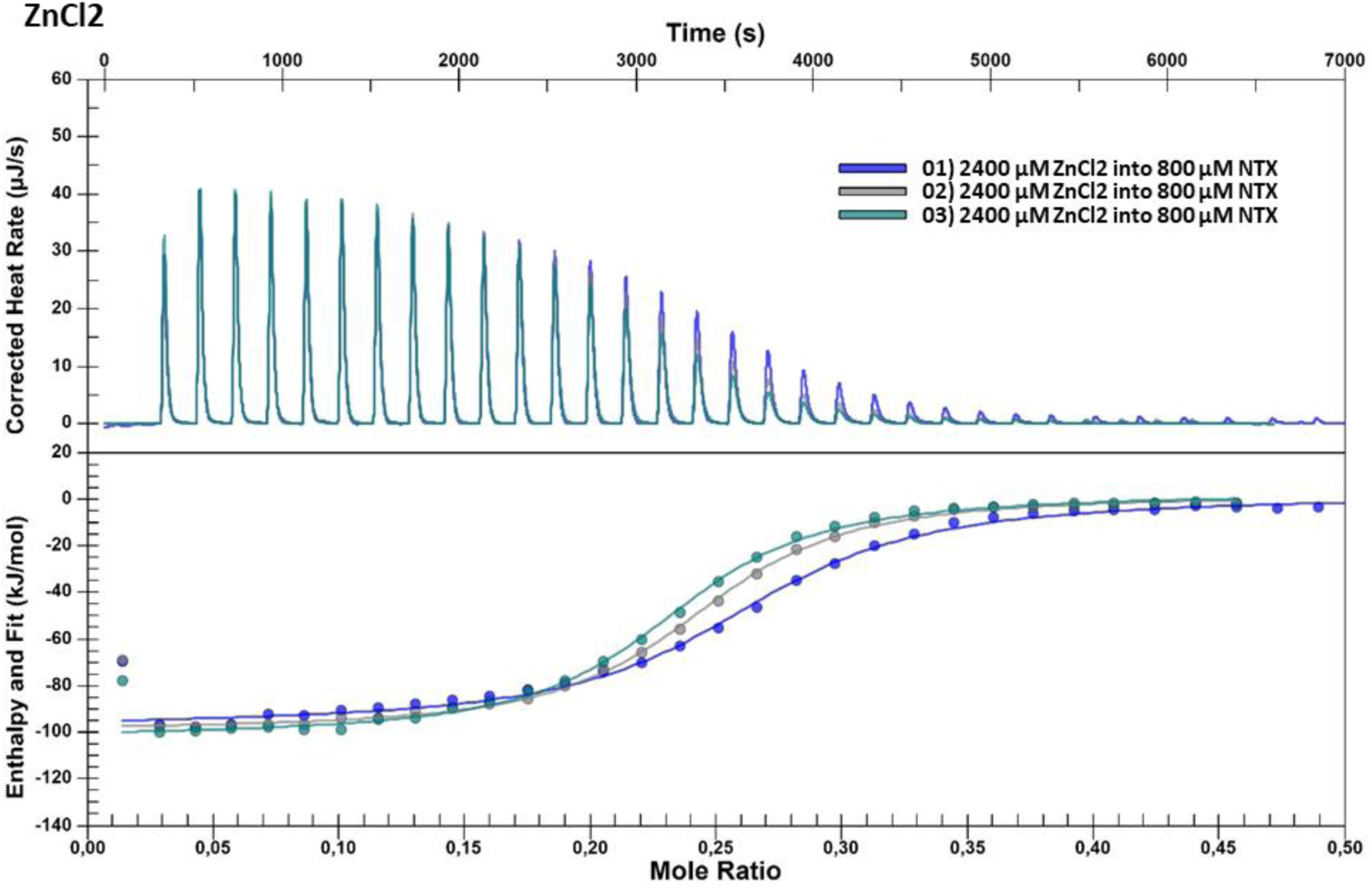
ITC results of NTX and ZnCl_2_. Performed in triplicates.

**Figure SI 3:**
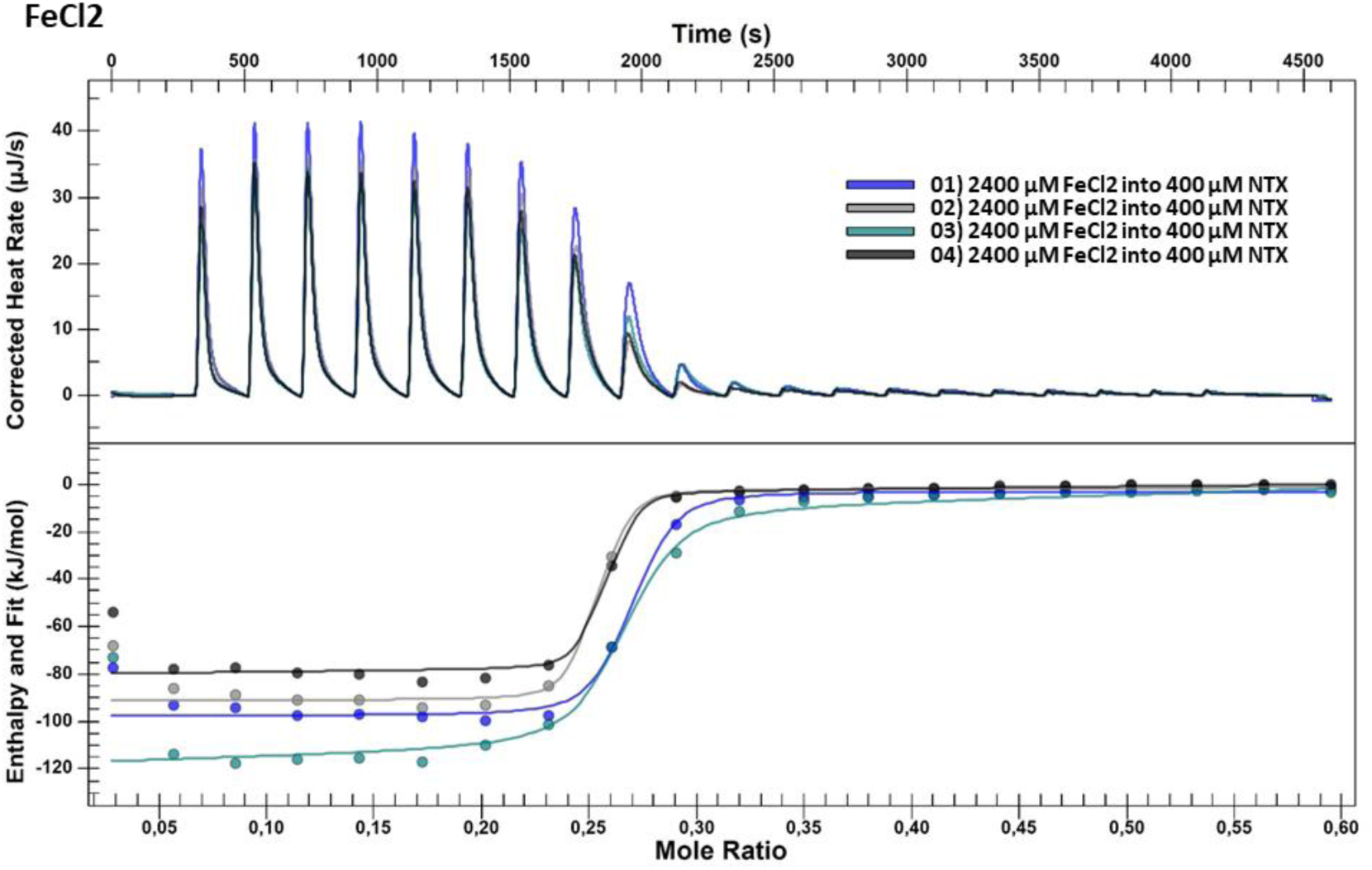
ITC results of NTX and FeCl_2_. Performed in quadruplicates.

**Figure SI 4:**
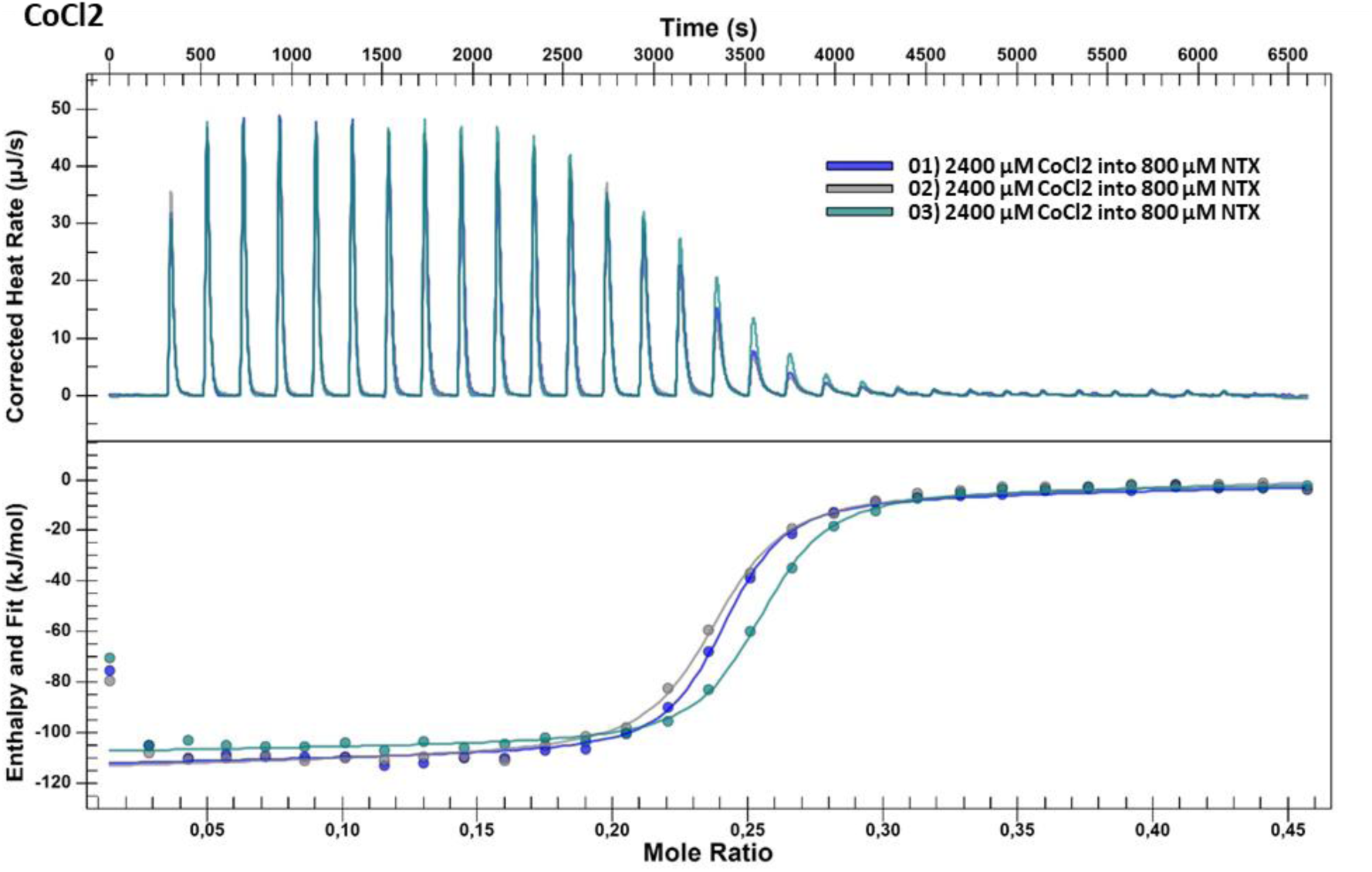
ITC results of NTX and CoCl_2_. Performed in triplicates.

**Figure SI 5:**
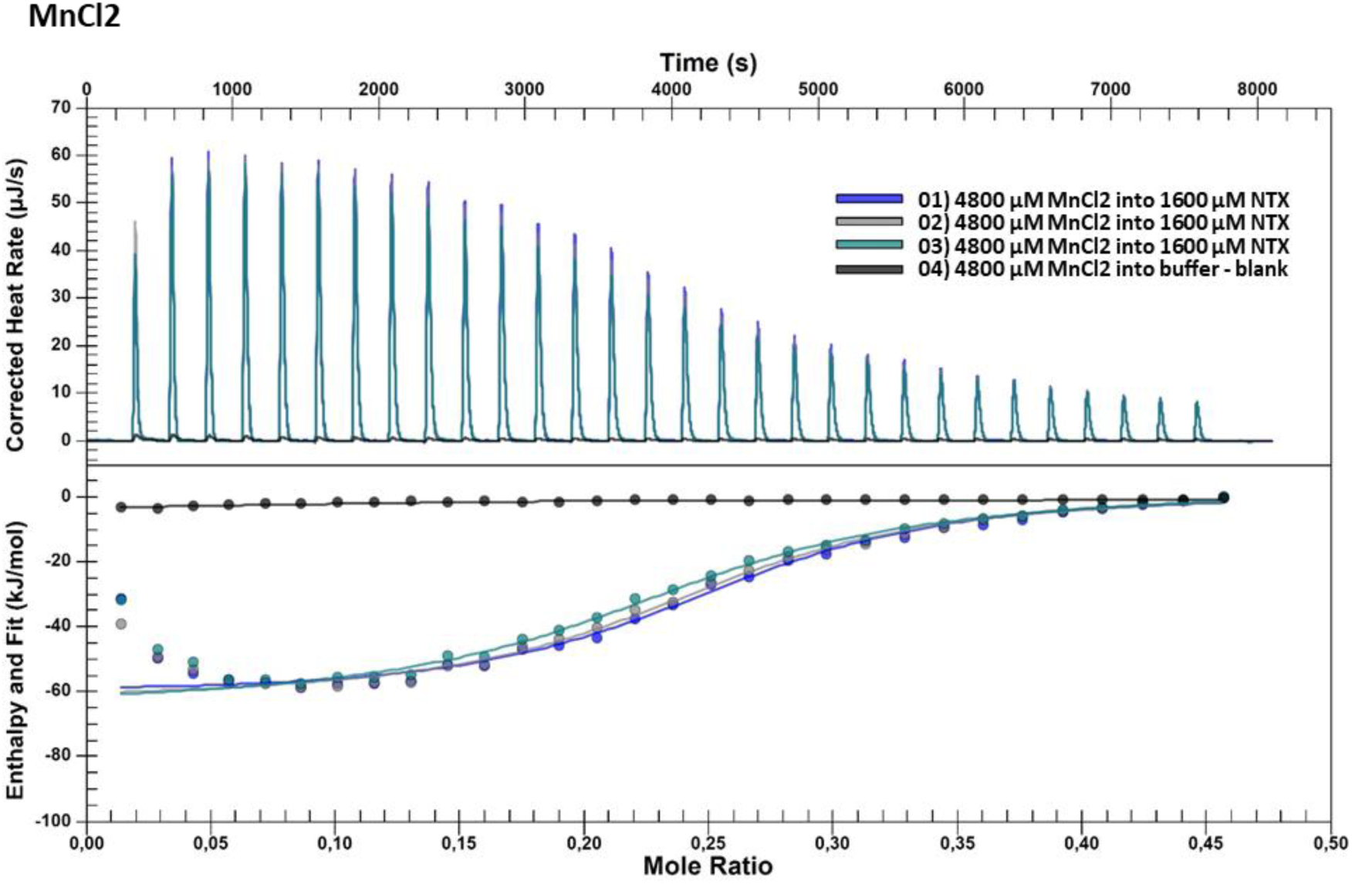
ITC results of NTX and MnCl_2_. Performed in triplicates with a blank titration into buffer, to check dilution heat.

**Figure SI 6:**
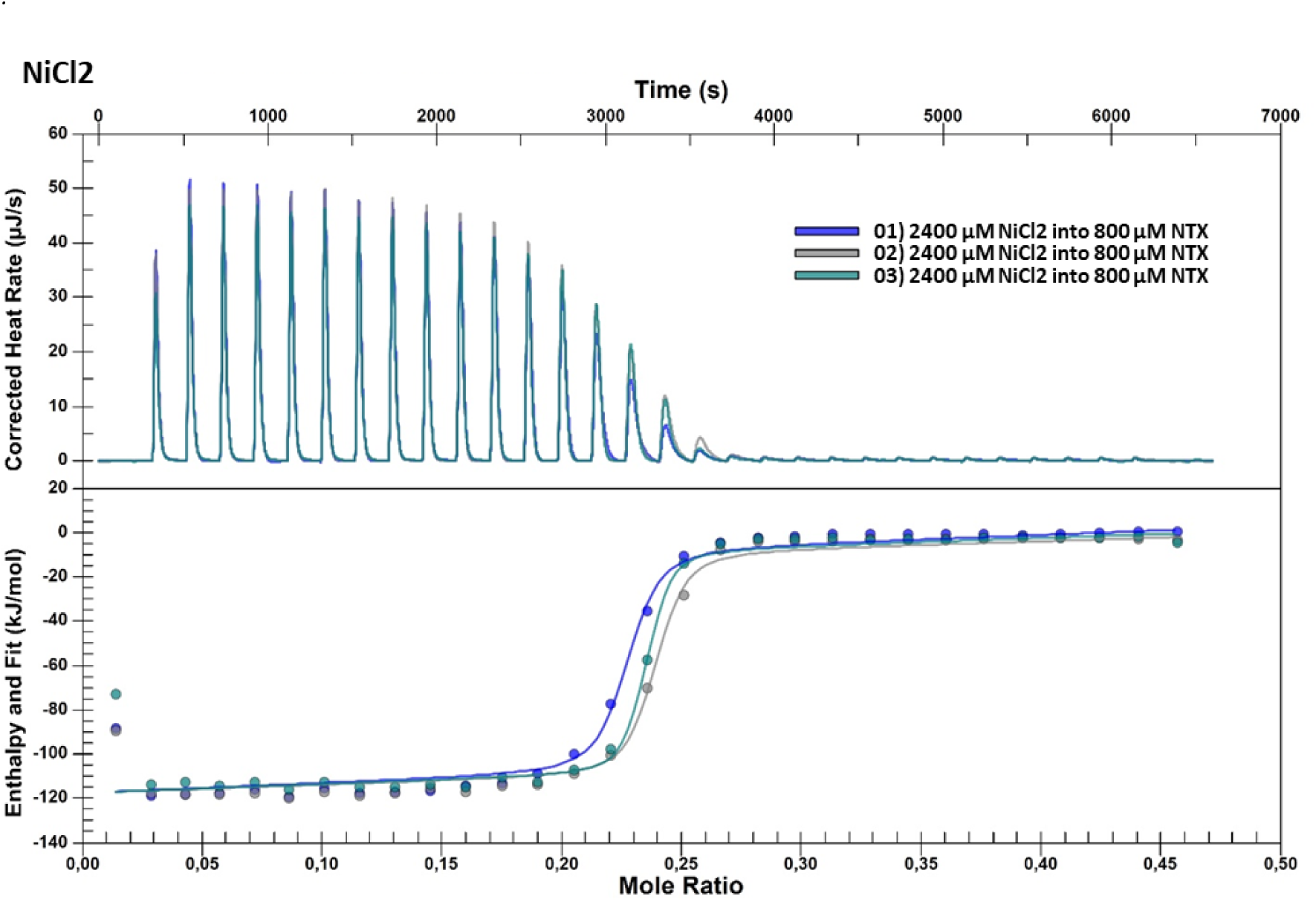
ITC results of NTX and NiCl_2_. Performed in triplicates.

**Figure SI 7:**
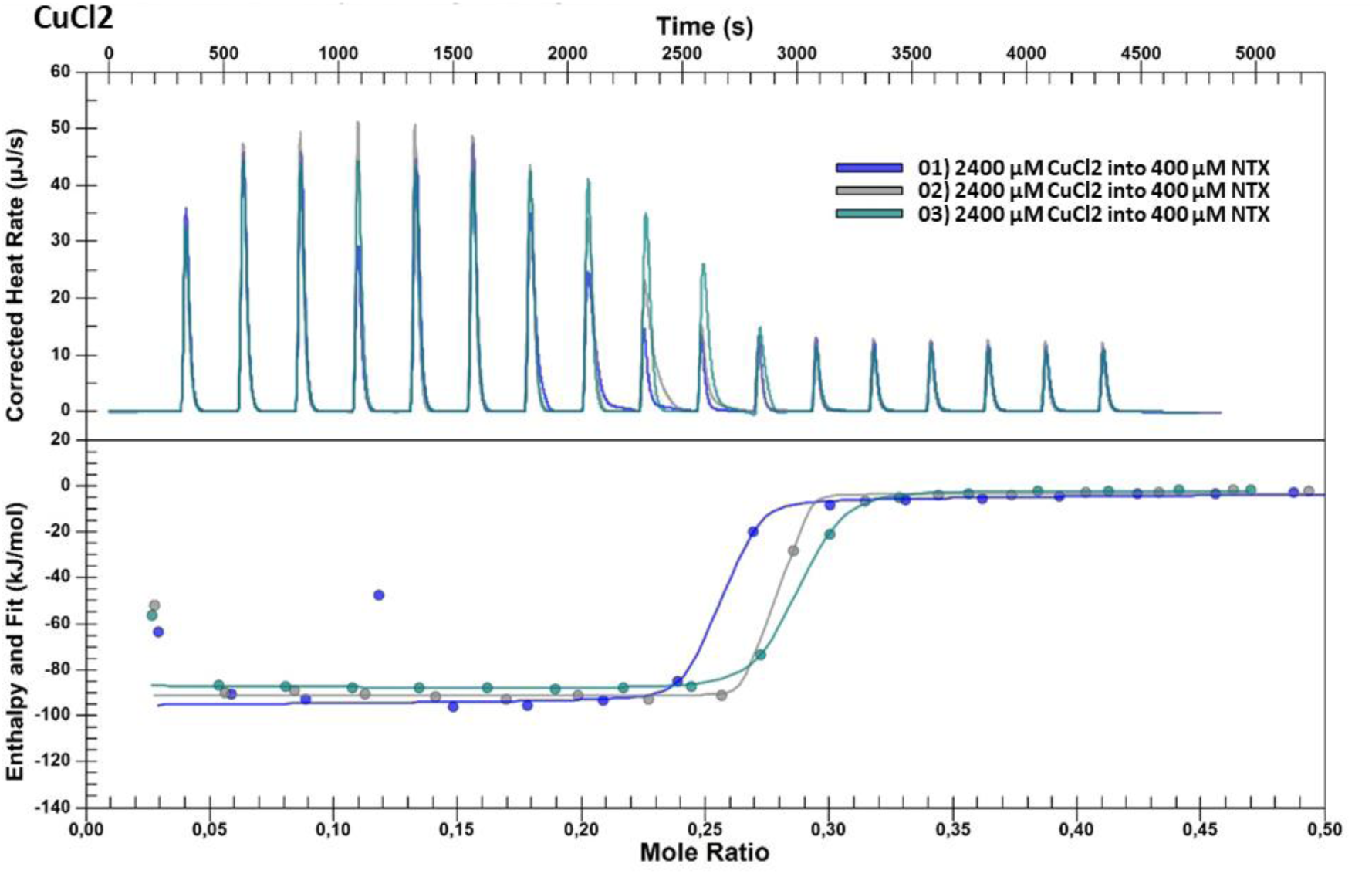
ITC results of NTX and CuCl_2_. Performed in triplicates.

**Figure SI 8:**
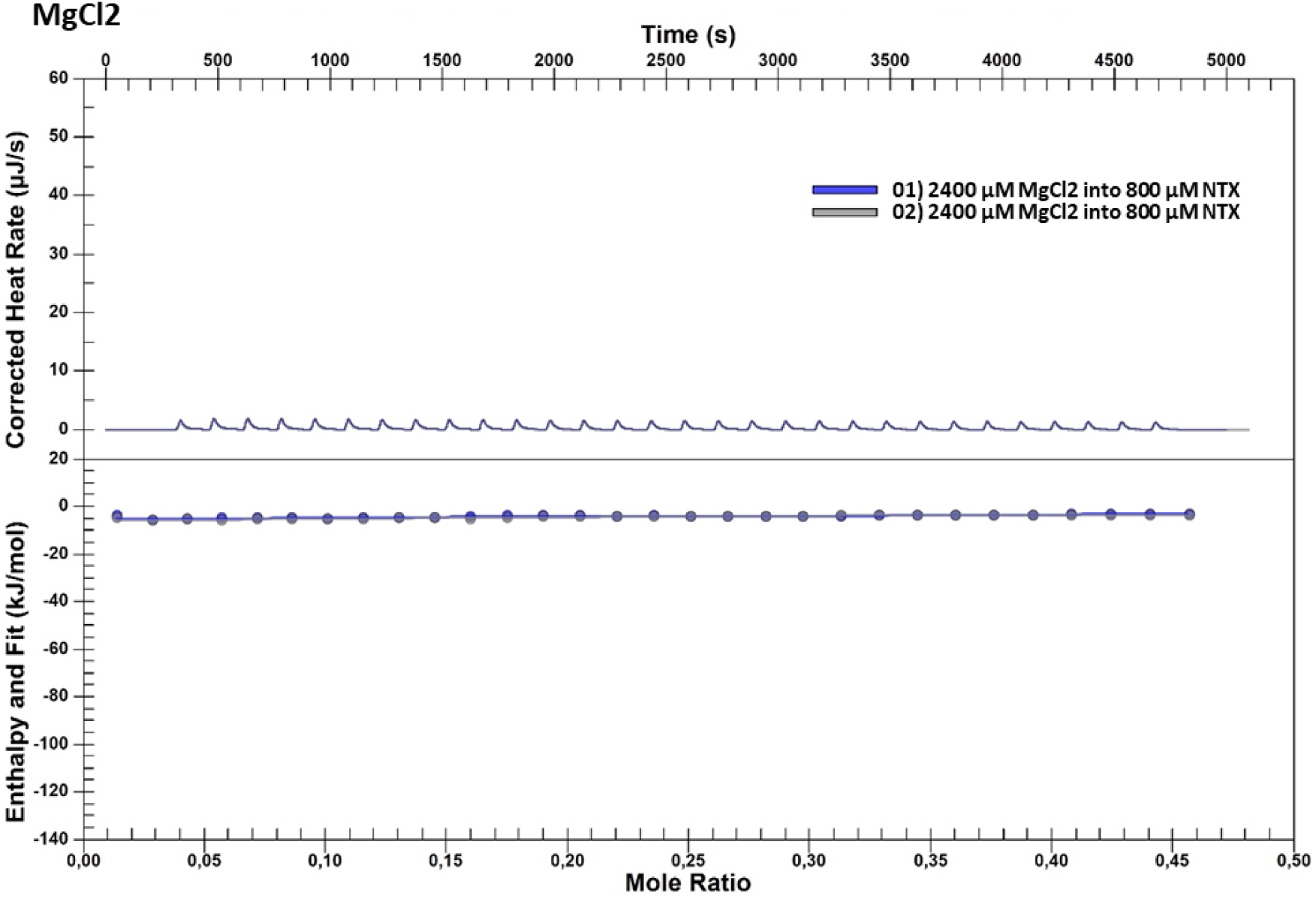
ITC results of NTX and MgCl_2_. Performed in duplicates.

**Figure SI 9:**
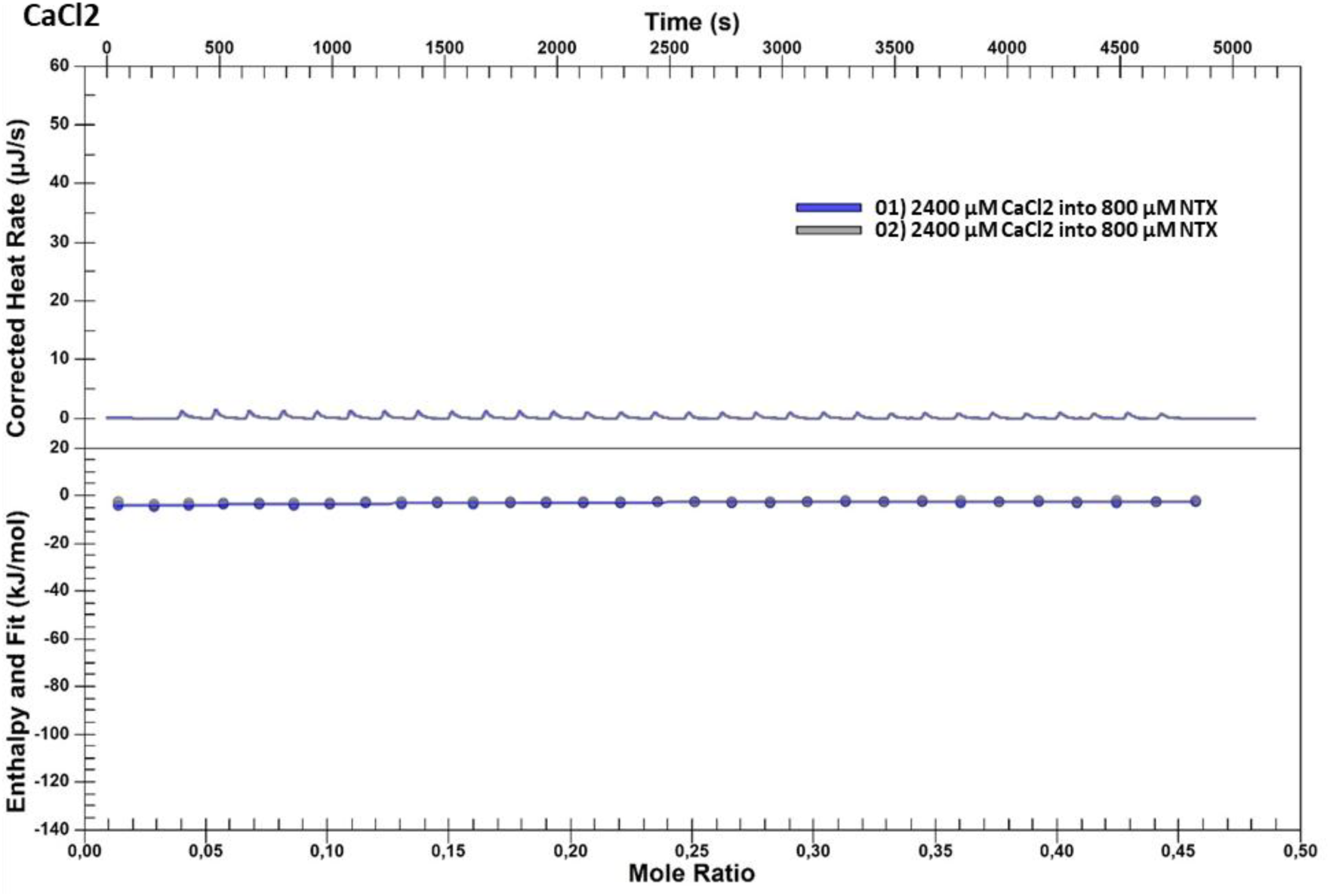
ITC results of NTX and CaCl_2_. Performed in duplicates.

**Figure SI 10:**
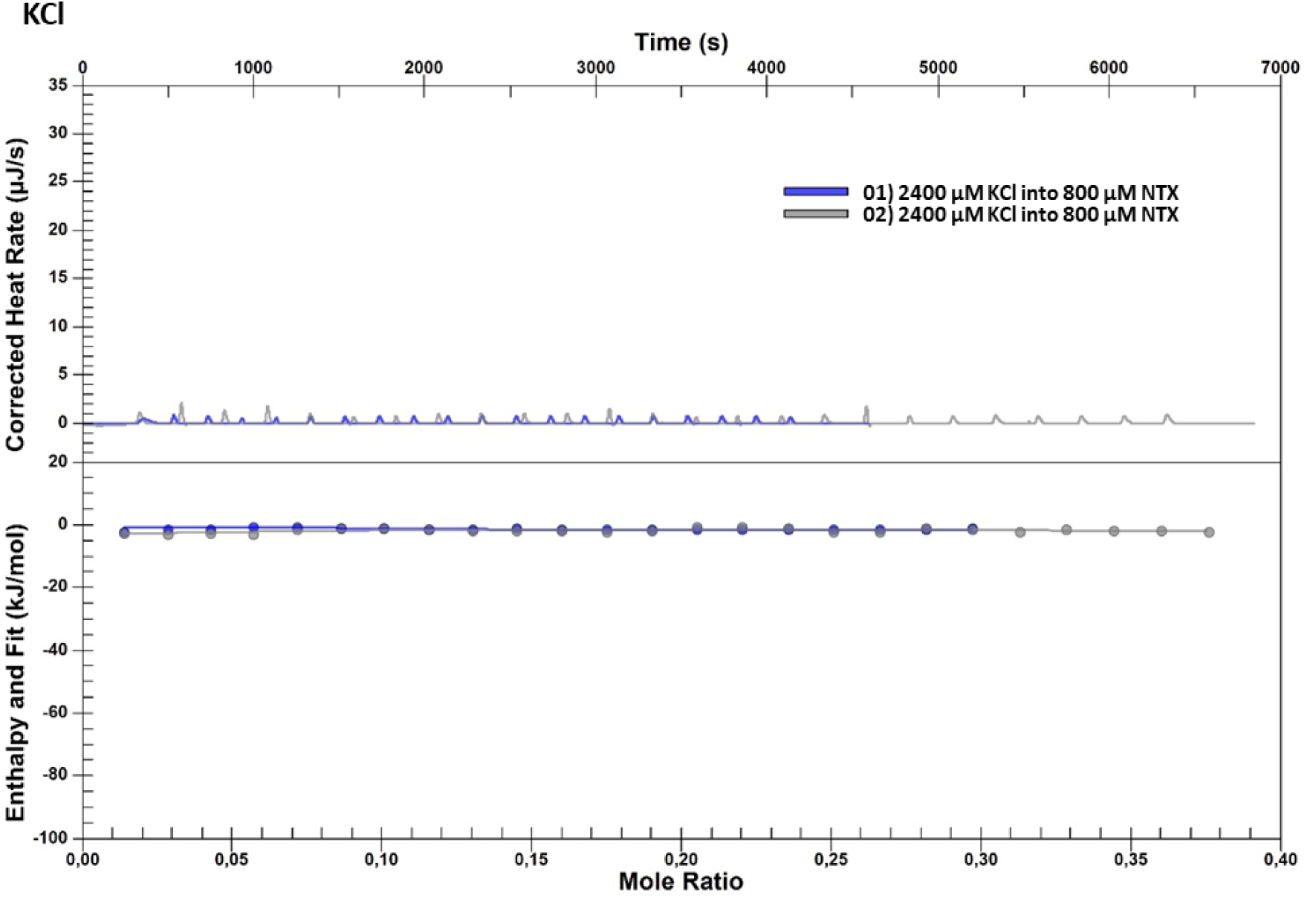
ITC results of NTX and KCl. Performed in duplicates.

**Figure SI 11:**
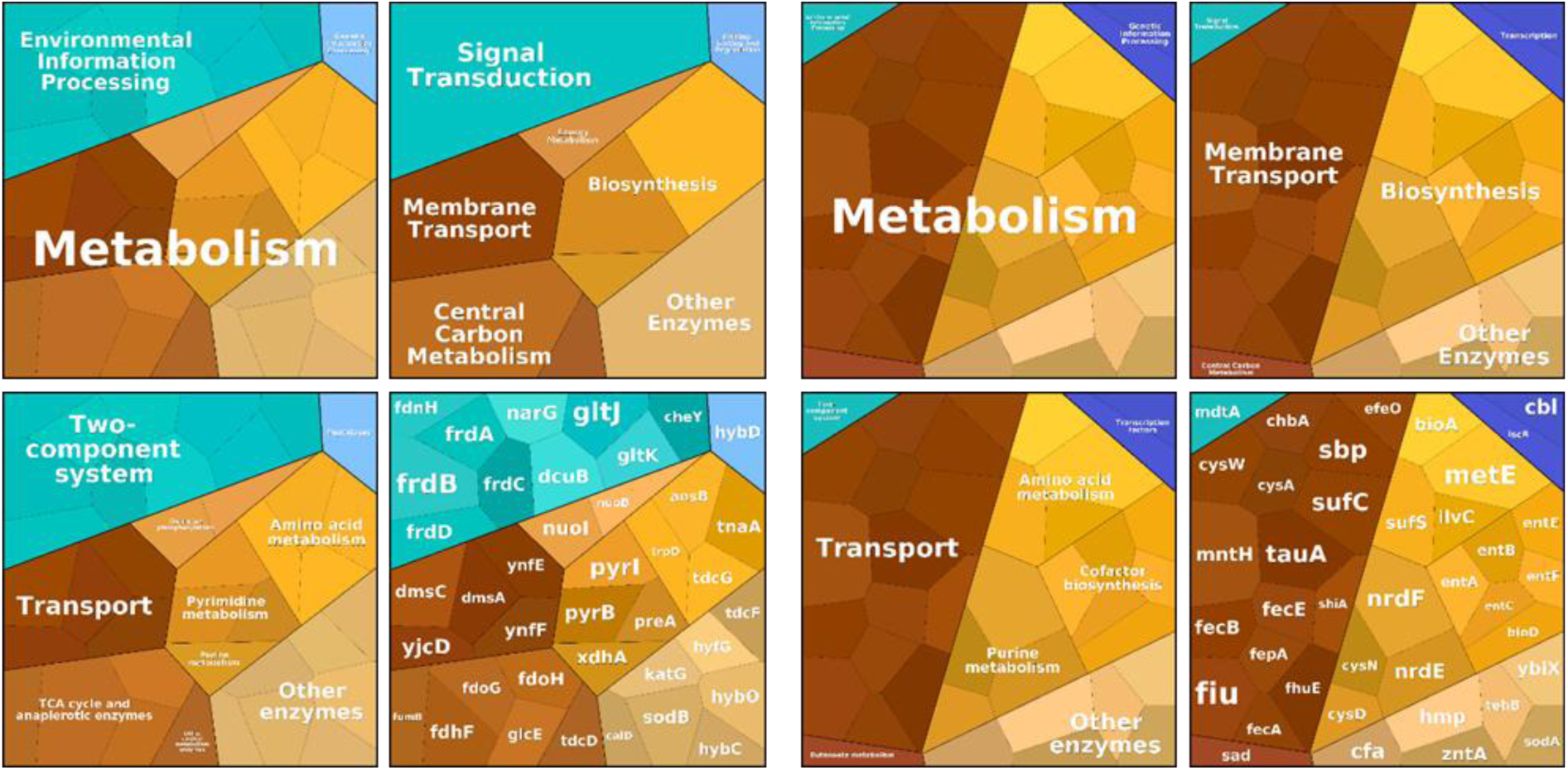
KEGG pathway analysis (Voronoi plots (Liebermeister et al, 2014)) shows clear influence of NTX treatment on metabolism of E. coli. Left: Voronoi plot of downregulated proteins (log2 < -2.5); right: Voronoi plot of upregulated proteins (log2 > 1.5). Size of the respective field is based on differential expression (log2 value), while colours correspond to KEGG annotation. Non-mapped proteins are excluded and shown in **Table SI 1**.

**Table SI 1:** Proteins missing from Voronoi plots. Annotations are not matched with KEGG reference resulting in loss of respective protein from Voronoi plot.

| KEGG ID | Protein | Description | Log<sub>2</sub> difference |
| --- | --- | --- | --- |
| b2713 | HydN | Electron transport protein | -4.38 |
| b2721 | HycE | Formate hydrogenlyase subunit 5 | -4.32 |
| b2727 | HypB | Hydrogenase maturation factor | -3.49 |
| b2214 | ApbE | FAD:protein FMN transferase | -3.41 |
| b1687 | YdiJ | D-2-hydroxyglutarate dehydrogenase | -3.37 |
| b2719 | HycG | Formate hydrogenlyase | -3.21 |
| b1541 | YdfZ | Putative selenoprotein | -2.84 |
| b1256 | NuoI | NADH-quinone oxidoreductase | -2.58 |
| b1681 | SufD | FeS cluster assembly protein | 4.86 |
| b1683 | SufB | FeS cluster assembly protein | 3.56 |
| b1684 | SufA | FeS cluster assembly protein | 3.32 |
| b1679 | SufE | Cystein desulphuration protein | 3.07 |
| b0597 | EntH | Proofreading thioesterase | 2.95 |
| b4209 | YtfE | Iron-Sulphur cluster repair protein | 2.87 |
| b1287 | YciQ | Uncharacterised | 2.79 |
| b4473 | Smf | Protein Smf | 2.62 |
| b4292 | FecR | Protein FecR | 2.40 |
| b1340 | SmrA | Probable DNA endonuclease | 2.17 |
| b0281 | IntF | Prophage integrase | 1.91 |
| b1495 | YddB | Uncharacterised | 1.81 |
| b1020 | PhoH | Protein PhoH | 1.72 |
| b1452 | YncE | Uncharacterised | 1.68 |
| b2142 | YohK | Inner membrane protein | 1.56 |

**Figure SI 12:**
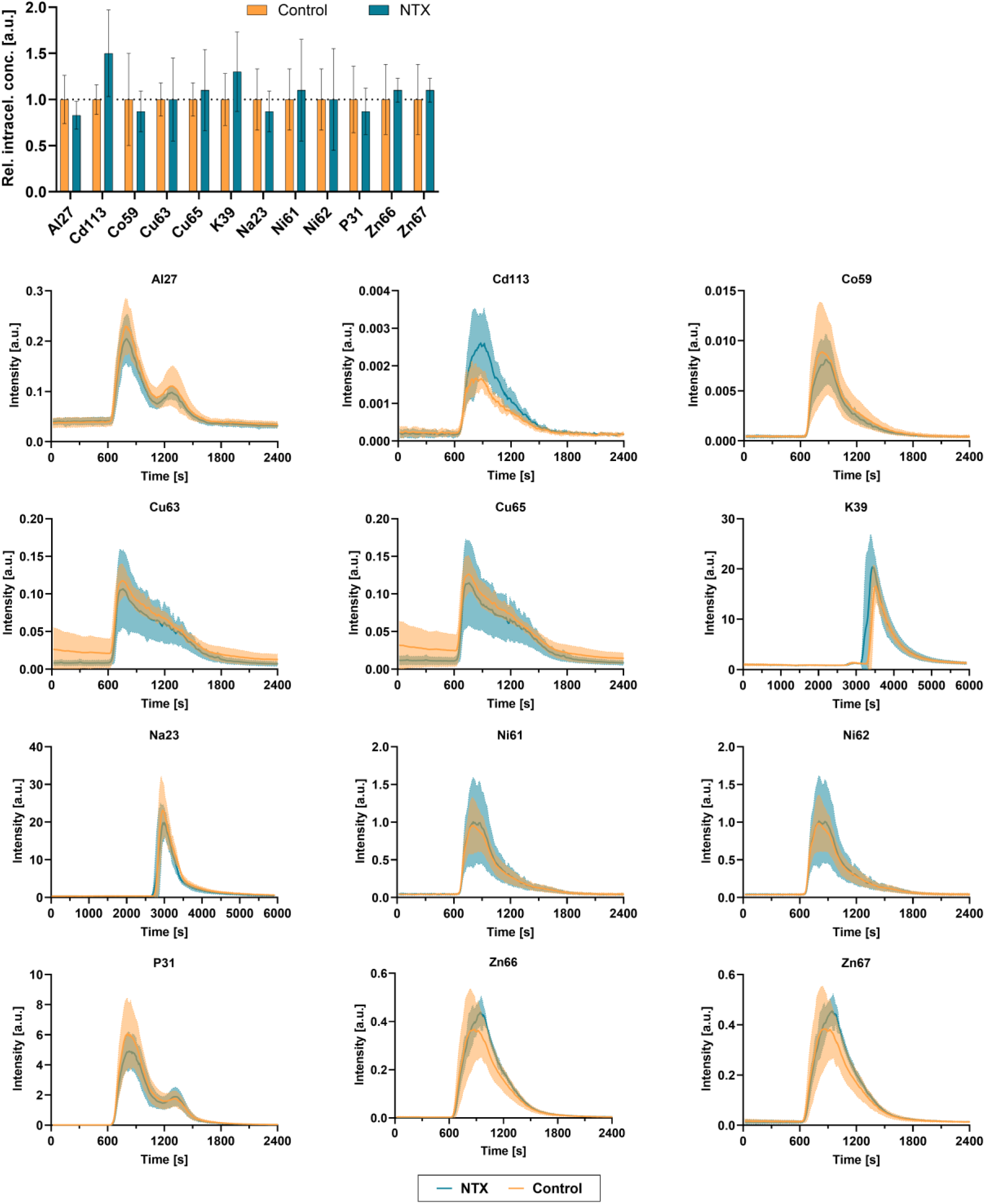
SEC-ICP-MS of non-significantly changed elements of NTX-treated E. coli.

**Scheme SI 1:**
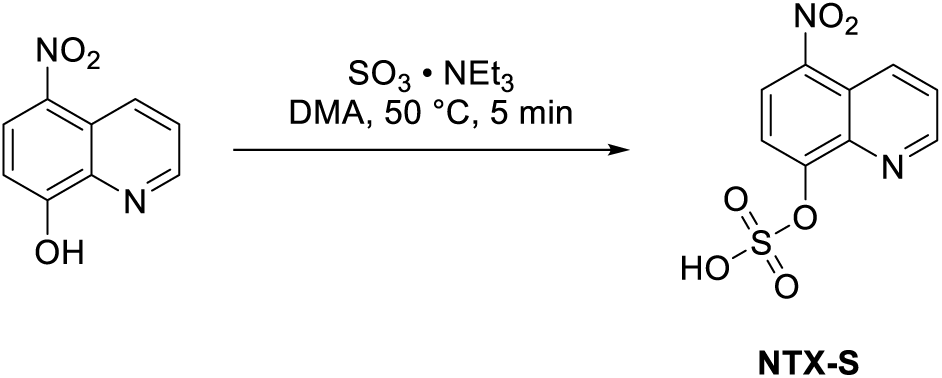
Chemical synthesis of NTX-S with total yield of 78.0%.

**Scheme SI 2:**
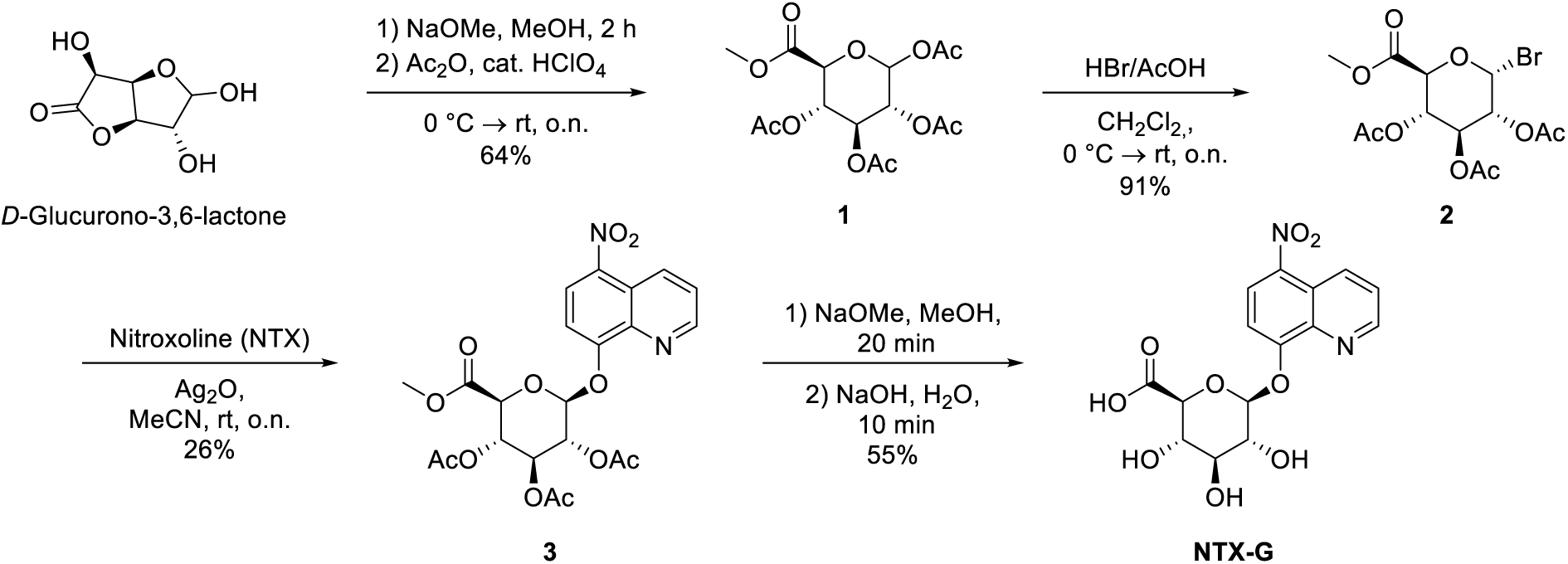
Chemical synthesis of NTX-G with total yield of 8.3%.

**Table SI 2:**
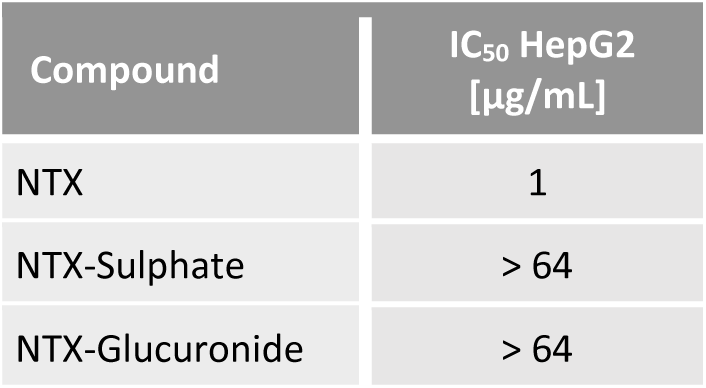
NTX metabolites are not cytotoxic against human liver cells.

**Figure SI 13:**
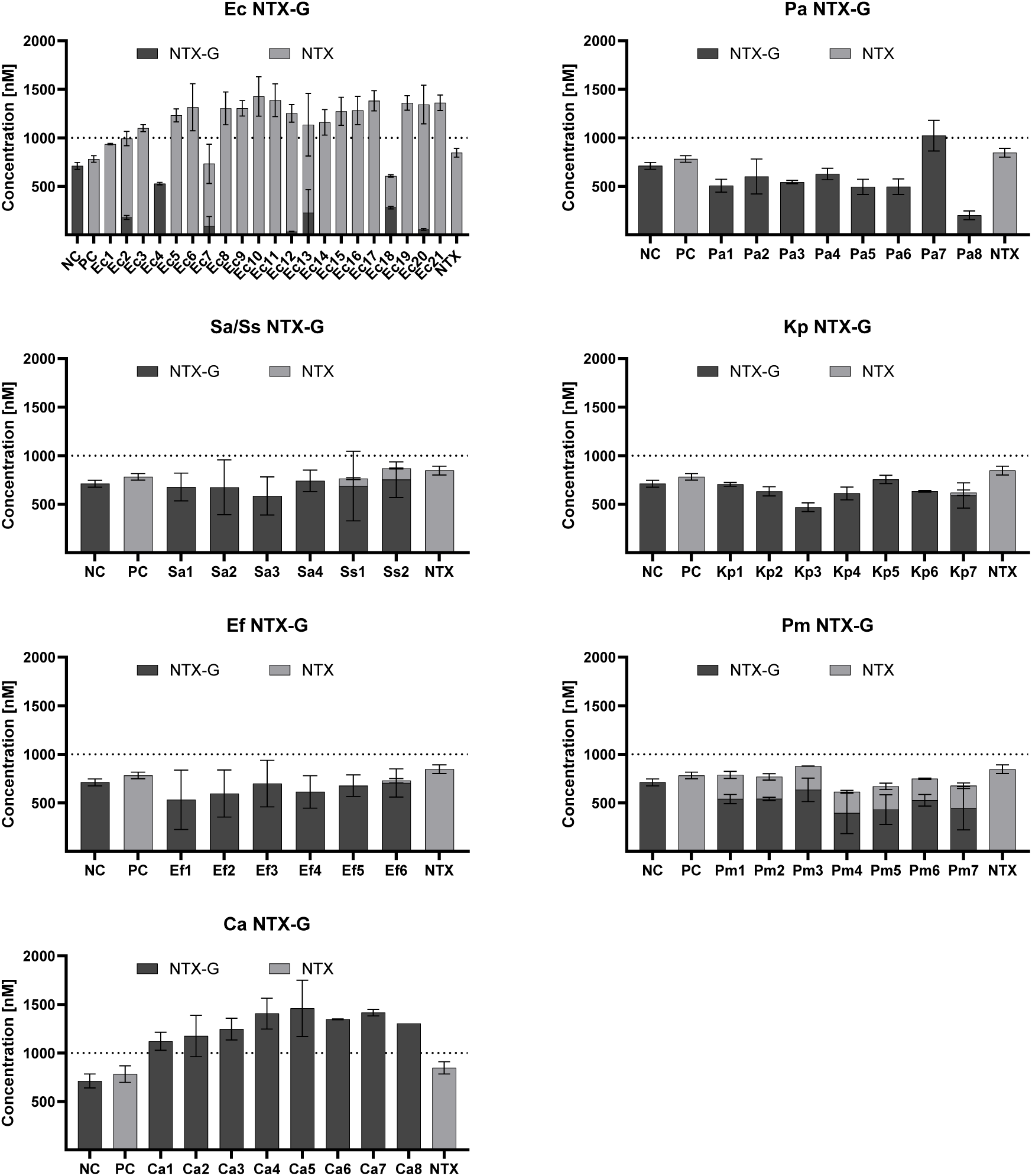
Conversion of NTX-G to NTX in presence of different clinical isolates displayed individually. Ec: E. coli, Pa: P. aeruginosa, Sa: S. aureus, Ss: S. saprophyticus, Kp: K. pneumoniae, Ef: E. faecalis, Pm: P. mirabilis, Ca: C. albicans. Experiment was done twice with mean and SEM displayed. E. coli ATCC25922 served as positive control (PC), medium w/o bacteria was used as negative control (NC). For reference, 1 µM NTX was displayed.

**Figure SI 14:**
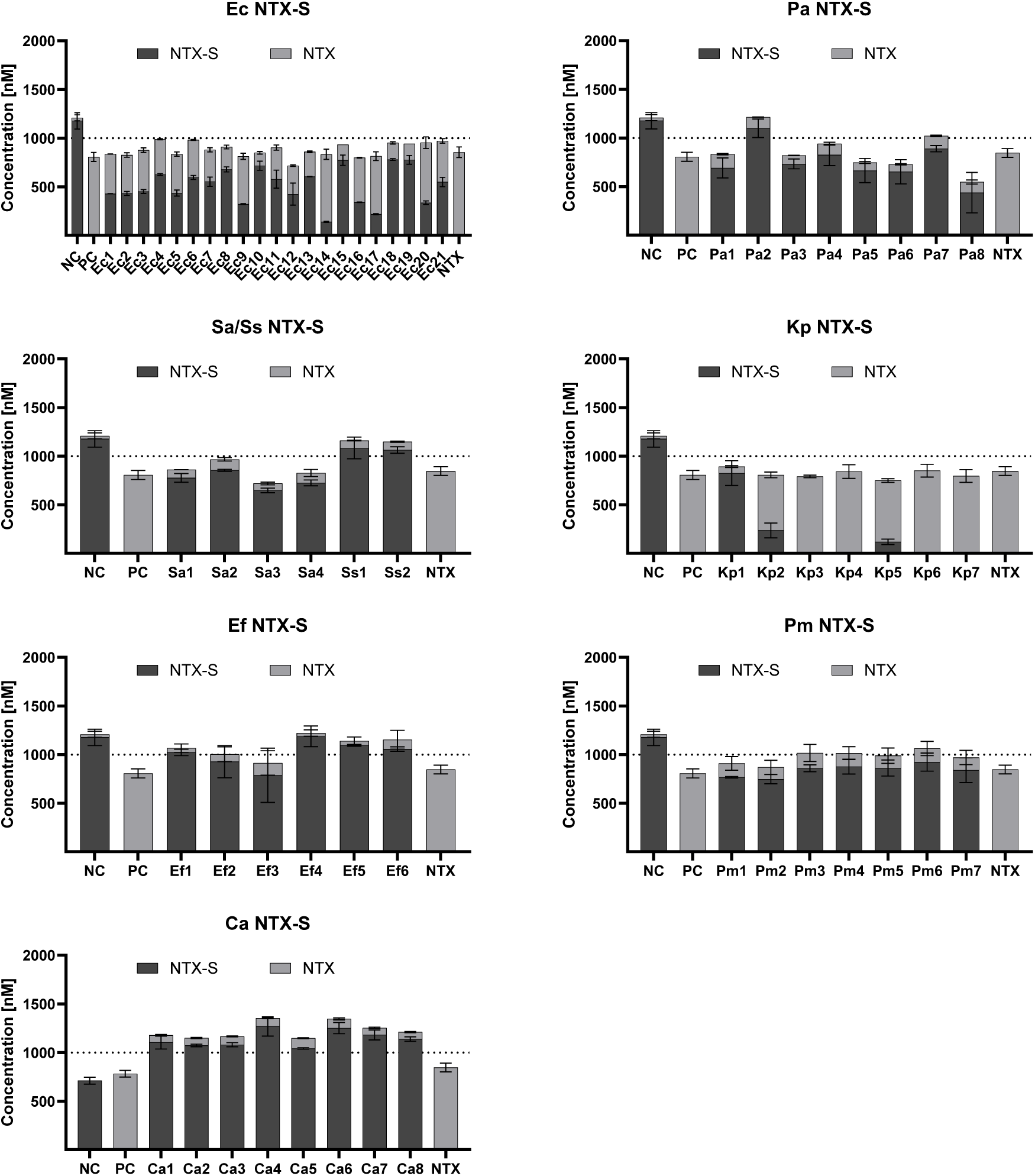
Conversion of NTX-S to NTX in presence of different clinical isolates displayed individually. Ec: E. coli, Pa: P. aeruginosa, Sa: S. aureus, Ss: S. saprophyticus, Kp: K. pneumoniae, Ef: E. faecalis, Pm: P. mirabilis, Ca: C. albicans. Experiment was done twice with mean and SEM displayed. E. coli ATCC25922 served as positive control (PC), medium w/o bacteria was used as negative control (NC). For reference, 1 µM NTX was displayed.

**Figure SI 15:**
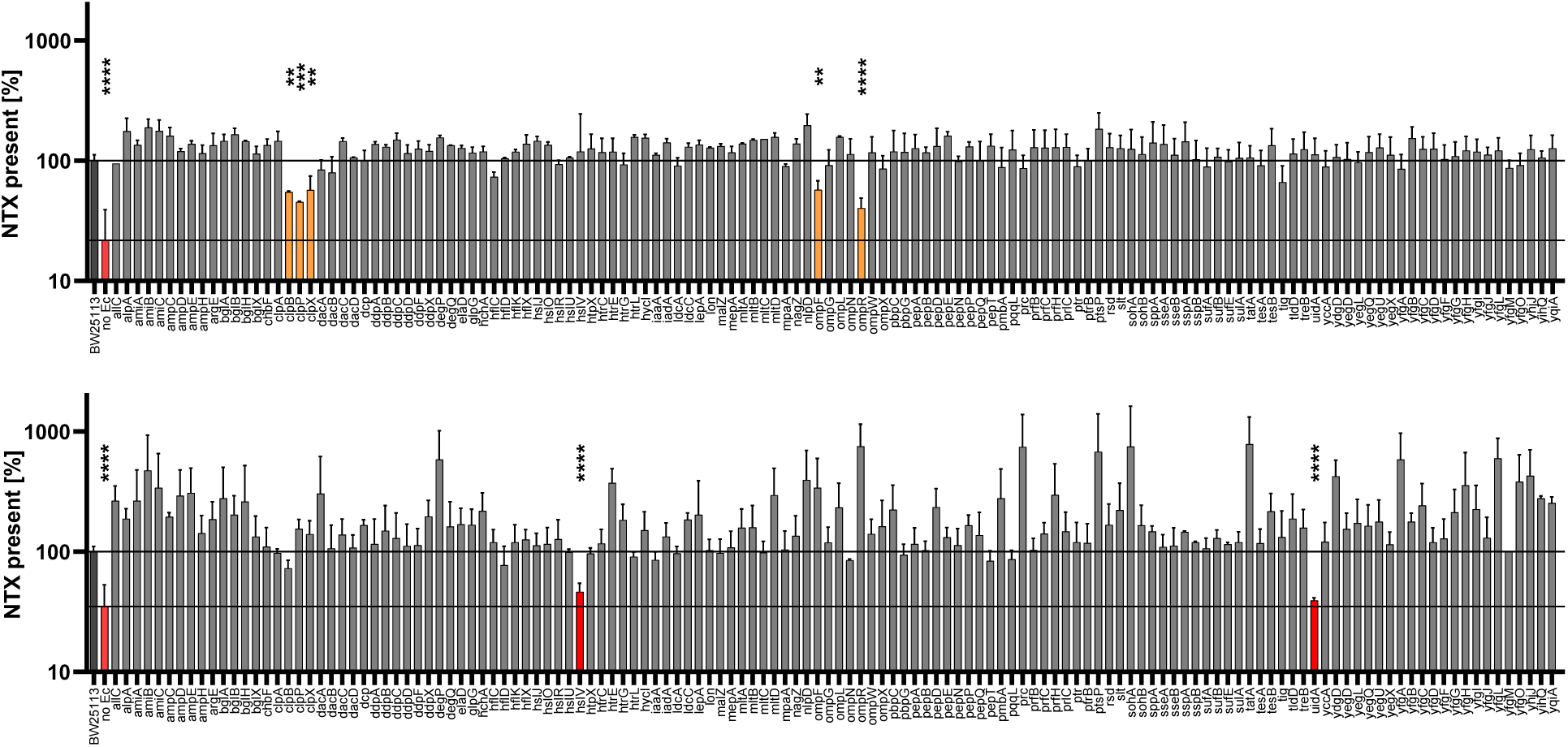
KEIO library screen indicates E. coli strains unable to convert NTX-metabolites into NTX. E. coli transposon strains (n = 134) with knockouts in different hydrolases etc were selected and screened for their NTX-conversion capability. The experiment was performed in pooled human urine (n = 2); the amount of NTX in each sample was measured, normalized first against the respective internal standard and then normalized against the library wildtype E. coli BW25113 set to 100%. A medium control without bacteria (no Ec) was used to determine baseline NTX values. Strains that produced significantly (t-test) less NTX compared to wildtype were coloured.

**Figure SI 16:**
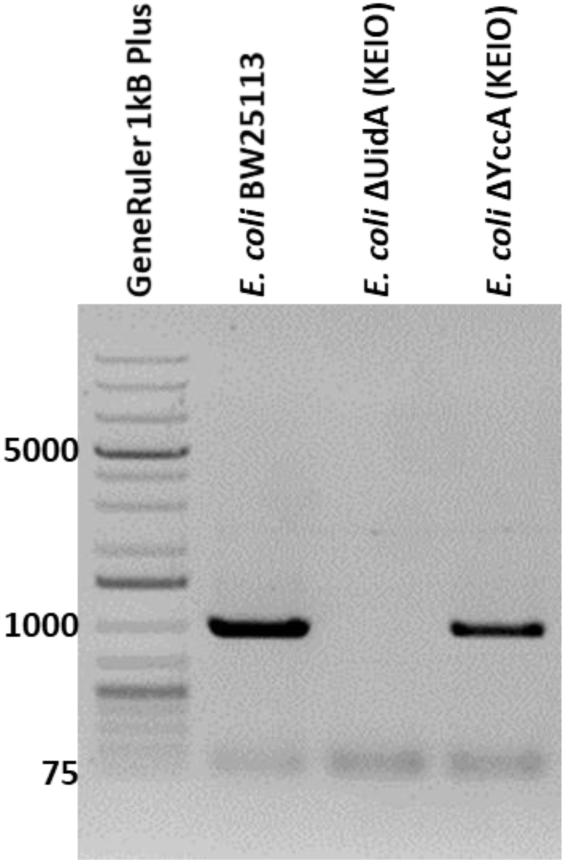
KEIO strain E. coli ΔuidA confirmed to not carry the uidA gene. Both KEIO wildtype E. coli BW25113 and unrelated KEIO strain ΔYccA were used as controls. Bands were amplified using a direct colony PCR with expected uidA band size 999 bps.

**Figure SI 17:**
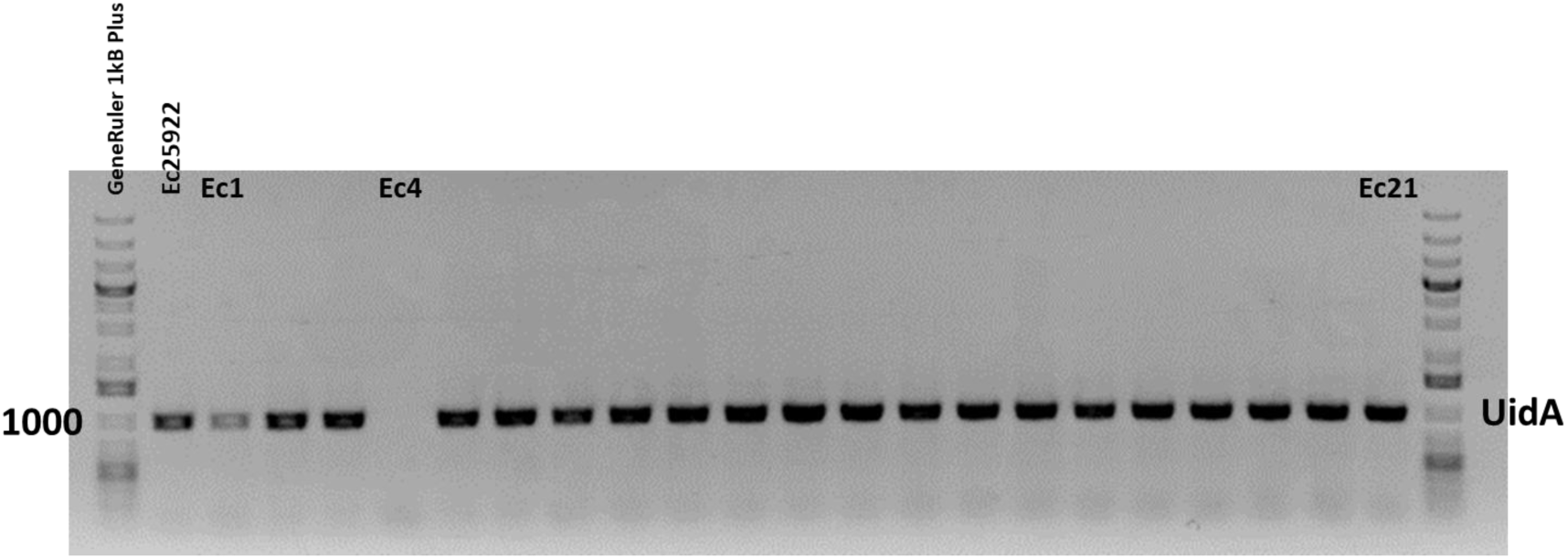
E. coli isolate 4 (Ec4) does not contain the uidA gene. Direct colony PCR with gene specific primers (UidA_F1 and UidA_R1) should result in a 999bp band if uidA is present.

**Figure SI 18:**
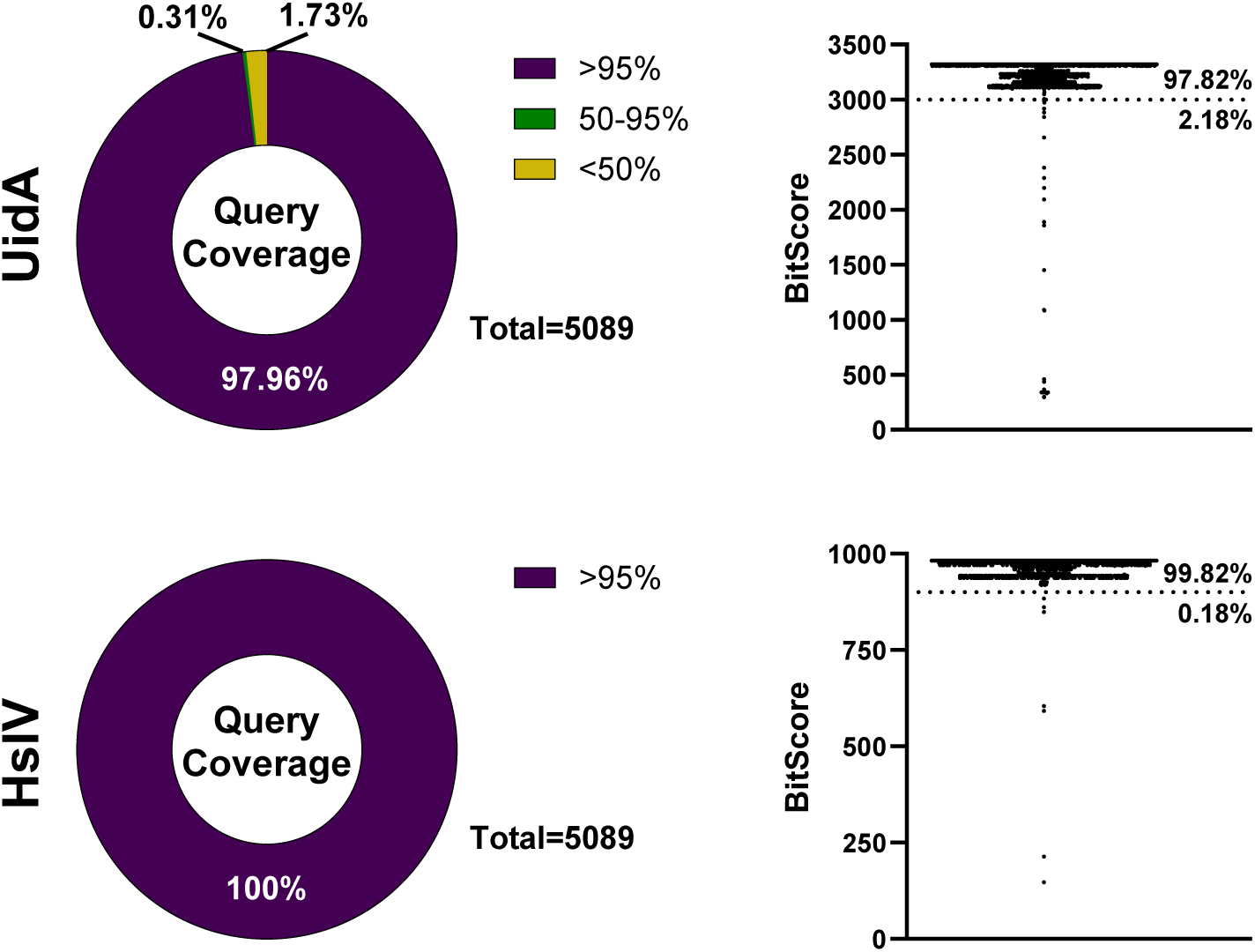
UidA and HslV belong to the core genome of E. coli. Genomes (n = 5089, GenBank annotated) were retrieved from NCBI and used as blast reference database for the uidA and hslV gene sequences performed with Geneious Prime Version 2025.2.2 (megablast). Left) Query coverage of the respective gene sequence against all 5089 strains. Right) Blast result (bitscore) plotted for every individual strain, separated according to quality of alignment.

## SI Methods

### General information for the synthesis of compounds

All air and moisture sensitive reactions were carried out in dried glassware (> 100 °C) under nitrogen atmosphere. Commercial chemicals and solvents were used without further purification. Anhydrous solvents were stored under nitrogen atmosphere and dried over molecular sieves. The products were purified by column chromatography on silica gel columns (Machery-Nagel 60, 0.063–0.2 mm) on a *Büchi Pure C-810 Flash* system. For reverse-phase chromatography (indicated by C_18_-SiO_2_), the *Flash* system was used with *Chromabond Flash RS 4* or *15 C_18_ec* columns and MeCN/H_2_O solvents. Analytical TLC was performed on pre-coated silica gel plates (Machery-Nagel, Alugram Xtra Sil G/UV_254_). Detection was accomplished with UV light (254 nm), KMnO_4_ solution or cerium(IV)/ ammonium molybdate solution. ^1^H and ^13^C NMR spectra were recorded on a Bruker Advance I 500 MHz spectrometer [^1^H 500 MHz and ^13^C 125 MHz]. Chemical shifts (δ) are reported in parts per million (ppm) relative to internal solvent signal. Peaks were assigned using (^1^H,^1^H)-COSY, (^1^H,^13^C)-HSQC and (^1^H,^13^C)-HMBC spectra. In all cases where no explicit diastereoselectivity is stated, the diastereoselectivity exceeds the ratio of at least 99:1 as observed in the crude ^1^H NMR analysis.

UPLC-MS measurement were performed on a Vanquish Flex UHPLC (Germering, Germany) system [before Dec 2023: on a Dionex (Germering, Germany) Ultimate 3000 RSLC system] equipped with Waters (Eschborn, Germany) BEH C_18_ column (100 × 2.1 mm, 1.7 μm) equipped with a Waters VanGuard BEH C_18_ 1.7 μm guard column. Separation of 1 µL sample was achieved by a linear gradient from (A) 0.1% FA in H_2_O to (B) 0.1% FA in ACN at a flow rate of 600 µL/min and 45 °C. The gradient was initiated by a 0.5 min isocratic step at 5% B, followed by an increase to 95% B in [insert gradient method here: 18 min/ 9 min/ 6 min] to end with a 2 min step at 95% B before re-equilibration with initial conditions. UV-vis spectra were recorded by a DAD in the range from 190 to 800 nm [for Dionex: 200-600 nm]. [insert MS method here]. Calibration was done automatically before every LC-MS run by injection of a basic sodium formate solution through a filled 20 µL loop switched into the LC flow at the beginning of each run. All MS analyses were acquired in the presence of the lock masses (C_12_H_19_F_12_N_3_O_6_P_3_, C_18_H_19_O_6_N_3_P_3_F_2_ and C_24_H_19_F_36_N_3_O_6_P_3_) which generate the [M+H]^+^ ions of 622.0289; 922.0098 and 1221.9906.

### MS methods

#### maXis MS-only

The LC flow was split to 75 μL/min before entering the Bruker Daltonics (Bremen, Germany) maXis 4G HR-qToF mass spectrometer equipped with an Apollo II ESI source. The split was set up with fused silica capillaries of 75 and 100 µm I.D. and a low dead volume tee junction (Upchurch). Mass spectra were acquired in centroid mode ranging from 150-2500 *m*/*z* at a 2 Hz scan rate in centroid mode. Mass spectrometry source parameters were set to 500 V as end plate offset; +4000 V as capillary voltage; nebuliser gas pressure 1 bar; dry gas flow of 5 L/min and a dry temperature of 200 °C. Ion transfer and quadrupole settings were set to funnel RF 350 Vpp.; multipole RF 400 Vpp as transfer settings and ion energy of 5 eV as well as a low mass cut of 300 *m/z*.

#### maXis MS^2^

Full scan spectra were acquired at 2 Hz followed by MS^2^ spectra acquisition at a scan speed of 0.68 Hz. CID (collision-induced dissociation) energy varied linearly from 35, 45, to 60 eV with respect to the precursor *m/z* from 500, 1000, to 2000 *m/z*. The ion cooler was set to ramp collision energy (80−120% of the set value) and ion cooler RF from 700 to 1000 Vpp for every MS^2^ scan. The precursor list was evaluated every 2 s to assign the upcoming precursors, and precursors were moved to an exclusion list for 0.2 min after two spectra were measured (typical chromatographic peak width was 0.10−0.15 min). Minimum precursor intensity was set to 5000.

### Synthesis NTX-G from NTX (refer to Scheme SI 2)

#### Methyl 1,2,3,4-tetra-acetyl-*D*-glucopyranuronate (1)

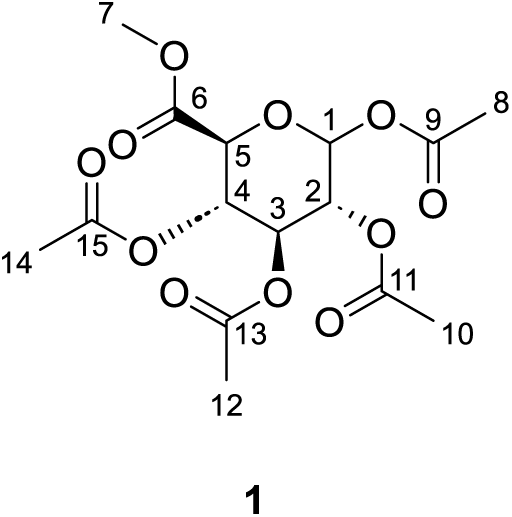

Compound 1 was synthesised with a modified protocol according to a published procedure (Herceg *et al*, 2018). 20.4 g (116 mmol, 1.0 eq.) D-(+)-glucuronic acid γ-lactone was suspended in anhydrous MeOH (115 mL, 1.0 M), cooled to 0 °C and 1.0 mL (5.40 mmol, 0.05 eq., 5.4 M in MeOH) NaOMe was added. The mixture was stirred for 2 h at room temperature before the solvent was removed under reduced pressure to obtain the crude as a brown syrup. In the next step, the crude was cooled to 0 °C and 93.0 mL (984 mmol, 8.5 eq.) Ac_2_O were added, followed by dropwise addition of 0.35 mL (5.79 mmol, 0.05 eq., 70%) HClO_4_. The mixture was allowed to warm to room temperature and stirred overnight. After filtration, the filtrate was poured onto ice, DCM was added and the mixture was neutralized with sat. aq. NaHCO_3_. The phases were separated, the aqueous phase was extracted with DCM (x3) and the combined organic phase was washed with brine. After drying over Na_2_SO_4_, the solvent was removed under reduced pressure. The crude was recrystallized from hot MeOH and compound 1 (28.0 g, 74.3 mmol, 64%) was obtained as a white solid.

^1^H NMR (500 MHz, CDCl_3_): δ = 5.76 (d, *J* = 7.8 Hz, 1 H, 1-H), 5.30 (t, *J* = 9.2 Hz, 1 H, 3-H), 5.24 (dd, *J* = 9.6, 9.5 Hz, 1 H, 4-H), 5.14 (dd, *J* = 8.8, 8.0 Hz, 1 H, 2-H), 4.17 (d, *J* = 9.6 Hz, 1 H, 5-H), 3.74 (s, 3 H, 7-H), 2.11 (s, 3 H, 8-H), 2.01–2.05 (m, 9 H, 10-H, 12-H, 14-H) ppm. ^13^C NMR (125 MHz, CDCl_3_): δ = 170.1/169.6/169.3/169.0/166.9 (C-6/C-9/C-11/C-13/C-15), 91.5 (C-1), 73.1 (C-5), 72.0 (C-3), 70.3 (C-2), 69.1 (C-4), 53.2 (C-7), 20.8 (C-8), 20.9/20.7/20.6 (C-10/C-12/C-14) ppm. HRMS (CI): m/z calcd for [C_15_H_20_O_11_ + Na]^+^: 399.08978, found: 399.08860.

#### (2*R*,3*R*,4*S*,5*S*,6*S*)-2-Bromo-6-(methoxycarbonyl)tetrahydro-2*H*-pyran-3,4,5-triyl triacetate (2)

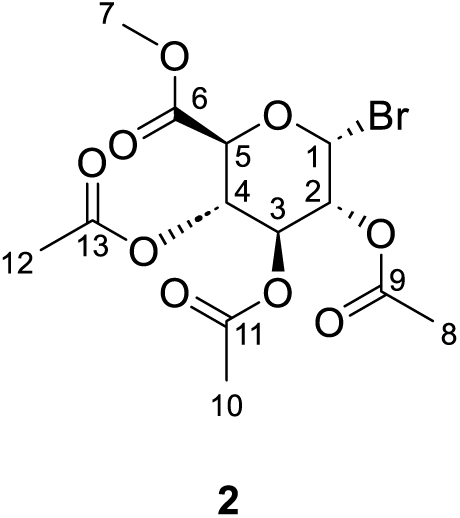

Compound 2 was synthesised with a modified protocol according to a published procedure (Walther *et al*, 2019). 5.03 g (15.1 mmol, 1.0 eq.) methyl 1,2,3,4-tetra-acetyl-D-glucopyranuronate (1) was dissolved in DCM (25 mL, 0.6 M) and cooled to 0 °C. 50.0 mL (836 mmol, 55.6 eq., 33w% in AcOH) HBr were added dropwise and the mixture was stirred for 5 h at room temperature. The mixture was poured onto ice and diluted with DCM. The aqueous phase was extracted with DCM (x3), the phases were separated and the organic phase was washed with cold H_2_O (x3), sat. aq. NaHCO_3_ (x3) and dried over Na_2_SO_4_. The solvent was removed under reduced pressure and the crude was dried *in vacuo*. Compound 2 (5.43 g, 13.7 mmol, 91%) was obtained as an off-white solid and used in the next step without further purification.

^1^H NMR (500 MHz, CDCl_3_): δ = 6.64 (d, *J* = 4.0 Hz, 1 H, 1-H), 5.61 (t, *J* = 9.8 Hz, 1 H, 3-H), 5.24 (tt, *J* = 9.9 Hz, 1 H, 4-H), 4.85 (dd, *J* = 10.0, 4.0 Hz, 1 H, 2-H), 4.58 (d, *J* = 9.6 Hz, 1 H, 5-H), 3.76 (s, 3 H, 7-H), 2.10 (s, 3 H, 8-H), 2.03–2.07 (m, 6 H, 8-H, 10-H) ppm. ^13^C NMR (125 MHz, CDCl_3_): δ = 169.8/169.8/169.6 (C-9/C-11/C-13), 166.8 (C-6), 85.5 (C-1), 72.2 (C-5), 70.5 (C-3), 69.4 (C-2), 68.6 (C-4), 53.3 (C-7), 20.8/20.8/20.6 (C-8/C-10/C-12) ppm. HRMS (CI): m/z calcd for [C_13_H_17_BrO_9_ + Na]^+^: 418.99482, found: 418.99313.

#### (2*S*,3*S*,4*S*,5*R*,6*S*)-2-(methoxycarbonyl)-6-((5-nitroquinolin-8-yl)oxy)tetrahydro-2*H*-pyran-3,4,5-triyl triacetate (3)

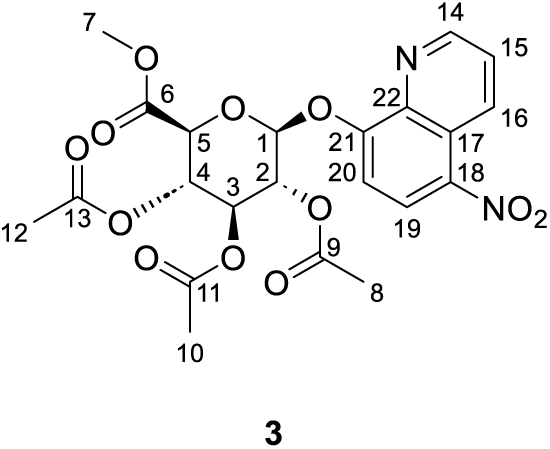

70.0 mg (368 μmol, 1.0 eq.) Nitroxoline (NTX, 5-nitroquinolin-8-ol) was dissolved in anhydrous MeCN (3.1 mL, 0.12 M), 292 mg (736 μmol, 2.0 eq.) (2*R*,3*R*,4*S*,5*S*,6*S*)-2-bromo-6-(methoxycarbonyl)tetra-hydro-2*H*-pyran-3,4,5-triyl triacetate (2) and 299 mg (1.29 mmol, 3.5 eq.) Ag_2_O were added and the mixture was stirred at room temperature overnight in the dark. The mixture was filtered over celite (washed with MeCN and EtOAc) and the solvent was removed under reduced pressure. The crude was first purified by column chromatography (SiO_2_, *n*-hexane/EtOAc) followed by a second purification by RP-MPLC (C_18_-SiO_2_, MeCN/H_2_O). Compound 3 (48 mg, 95 μmol, 26%) was obtained as a white solid after lyophilisation. The ratio of β:α anomer was determined as 97:3 by ^1^H-NMR analysis. R_f_ (3) = 0.36 (PE/EtOAc 1:1).

^1^H NMR (500 MHz, CDCl_3_): δ = 9.15 (dd, *J* = 8.9, 1.2 Hz, 1 H, 16-H), 9.02 (dd, *J* = 4.0, 1.4 Hz, 1 H, 14-H), 8.45 (d, *J* = 8.7 Hz, 1 H, 19-H), 7.69 (dd, *J* = 8.8, 4.1 Hz, 1 H, 15-H), 7.47 (d, *J* = 8.7 Hz, 0.98/1 H, 9-H [β-3]), 7.42 (dd, *J* = 8.8 Hz, 0.03/1H, 9-H [α-3]), 6.32 (d, *J* = 3.7 Hz, 0.03/1 H, 1-H [α-3]), 5.99 (t, *J* = 9.8 Hz, 0.03/1 H, 3-H [α-3]), 5.70 (d, *J* = 7.0 Hz, 01 H, 1-H]), 5.51 (t, *J* = 7.7 Hz, 1 H, 2-H), 5.40–5.48 (m, 2 H, 3-H, 4-H), 4.26 (d, *J* = 8.8 Hz, 1 H, 6-H), 3.68 (s, 3 H, 7-H), 2.06–2.10 (m, 9 H, 8-H, 10-H, 12-H) ppm. ^13^C NMR (125 MHz, CDCl_3_): δ = 170.2/169.7/169.5 (C-9/C-11/C-13), 167.0 (C-6), 156.1 (C-21), 149.7 (C-14), 140.2 (C-18), 138.0 (C-22), 134.9 (C-16), 127.3 (C-19), 124.7 (C-15), 123.2 (C-17), 114.4 (C-20), 99.8 (C-1), 73.2 (C-5), 71.2 (C-2), 71.1 (C-3), 68.8 (C-4), 53.2 (C-7), 21.0/20.9/20.7 (C-8/C-10/C-12) ppm. HRMS (CI): m/z calcd for [C_22_H_22_N_2_O_12_ + H]^+^: 507.12455, found: 507.12316.

#### (2*S*,3*S*,4*S*,5*R*,6*S*)-3,4,5-Trihydroxy-6-((5-nitroquinolin-8-yl)oxy)tetrahydro-2*H*-pyran-2-carboxylic acid (NTX-G)

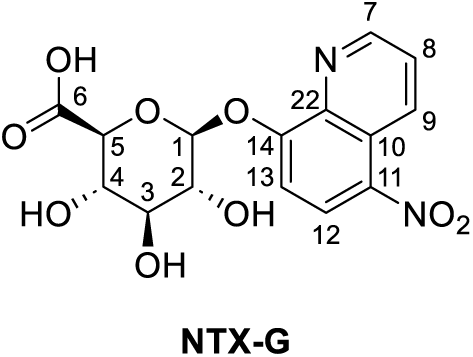

15.0 mg (30 μmol, 1.0 eq.) (2*S*,3*S*,4*S*,5*R*,6*S*)-2-(methoxycarbonyl)-6-((5-nitroquinolin-8-yl)oxy)tetra-hydro-2*H*-pyran-3,4,5-triyl triacetate (3) was dissolved in anhydrous MeOH (1.0 mL, 0.03 M), cooled to 0 °C and 5.4 μL (30 μmol, 1.0 eq., 5.4 M in MeOH) NaOMe was added. The mixture was stirred at room temperature for 20 minutes, cooled to 0 °C again and 30 μL (60 μmol, 2.0 eq., 2 M in H_2_O) NaOH was added. The mixture was stirred for another 10 minutes at room temperature, diluted with H_2_O and neutralized with weakly acidic ion exchanger resin. The solvent was removed by lyophilization and the crude was purified by RP-MPLC (C_18_-SiO_2_, MeCN/H_2_O + 0.1% formic acid). Compound NTX-G (6.0 mg, 16 μmol, 55%) was obtained as a white solid.

For further experiments, compound NTX-G was additionally purified by preparative HPLC to obtain 0.40 mg (indicated by name NTX-G-prep). The ratio of β:α anomer was determined as >95:5 by ^1^H-NMR analysis.

^1^H NMR (500 MHz, dmso-d_6_, compound NTX-G): δ = 9.05 (dd, *J* = 4.1, 1.6 Hz, 1 H, 7-H), 9.01 (dd, *J* = 8.8, 1.6 Hz, 1 H, 9-H), 8.54 (d, *J* = 8.8 Hz, 1 H, 12-H), 7.86 (dd, *J* = 8.8, 4.1 Hz, 1 H, 8-H), 7.54 (d, *J* = 8.8 Hz, 1 H, 13-H), 6.05 (d, *J* = 3.5 Hz, 0.06/1 H, 1-H [α-NTX-G]), 5.64 (d, *J* = 4.7 Hz, 1 H, 2-OH), 5.54 (d, *J* = 7.6 Hz, 0.95/1 H, 1-H [β-NTX-G]), 5.32 (bs, 1 H, 4-OH), 4.01 (d, *J* = 9.1 Hz, 1 H, 5-H), 3.49–3.54 (m, 1 H, 2-H), 3.37–3.45 (m, 2 H, 3-H, 4-H) ppm. ^1^H NMR (500 MHz, D_2_O, compound NTX-G-prep): δ = 9.08 (d, *J* = 8.7 Hz, 1 H, 9-H), 8.84 (d, *J* = 3.7 Hz, 1 H, 7-H), 8.43 (d, *J* = 8.7 Hz, 1 H, 12-H), 7.76 (dd, *J* = 8.6, 4.2 Hz, 1 H, 8-H), 7.40 (d, *J* = 8.7 Hz, 1 H, 13-H), 5.99 (d, *J* = 3.4 Hz, 0.04/1 H, 1-H [α-NTX-G]), 5.43 (d, *J* = 7.8 Hz, 0.95/1 H, 1-H [β-NTX-G]), 4.07 (d, *J* = 7.5 Hz, 1 H, 5-H), 3.85 (t, *J* = 8.2 Hz, 1 H, 2-H), 3.59–3.71 (m, 2 H, 3-H, 4-H) ppm. ^13^C NMR (125 MHz, D_2_O, compound NTX-G-prep): δ = 174.3 (C-6), 156.1 (C-14), 149.8 (C-7), 138.7 (C-11), 136.8 (C-22), 134.3 (C-9), 128.0 (C-12), 125.1 (C-8), 122.6 (C-10), 110.4 (C-13), 99.7 (C-1), 76.0 (C-5), 74.9 (C-3), 72.4 (C-2), 71.5 (C-4) ppm. HRMS (CI): m/z calcd for [C_15_H_14_N_2_O_9_ + H]^+^: 367.07721, found: 367.07633.

### Methyl 1,2,3,4-tetra-acetyl-D-glucopyranuronate (1) (Herceg *et al*, 2018)

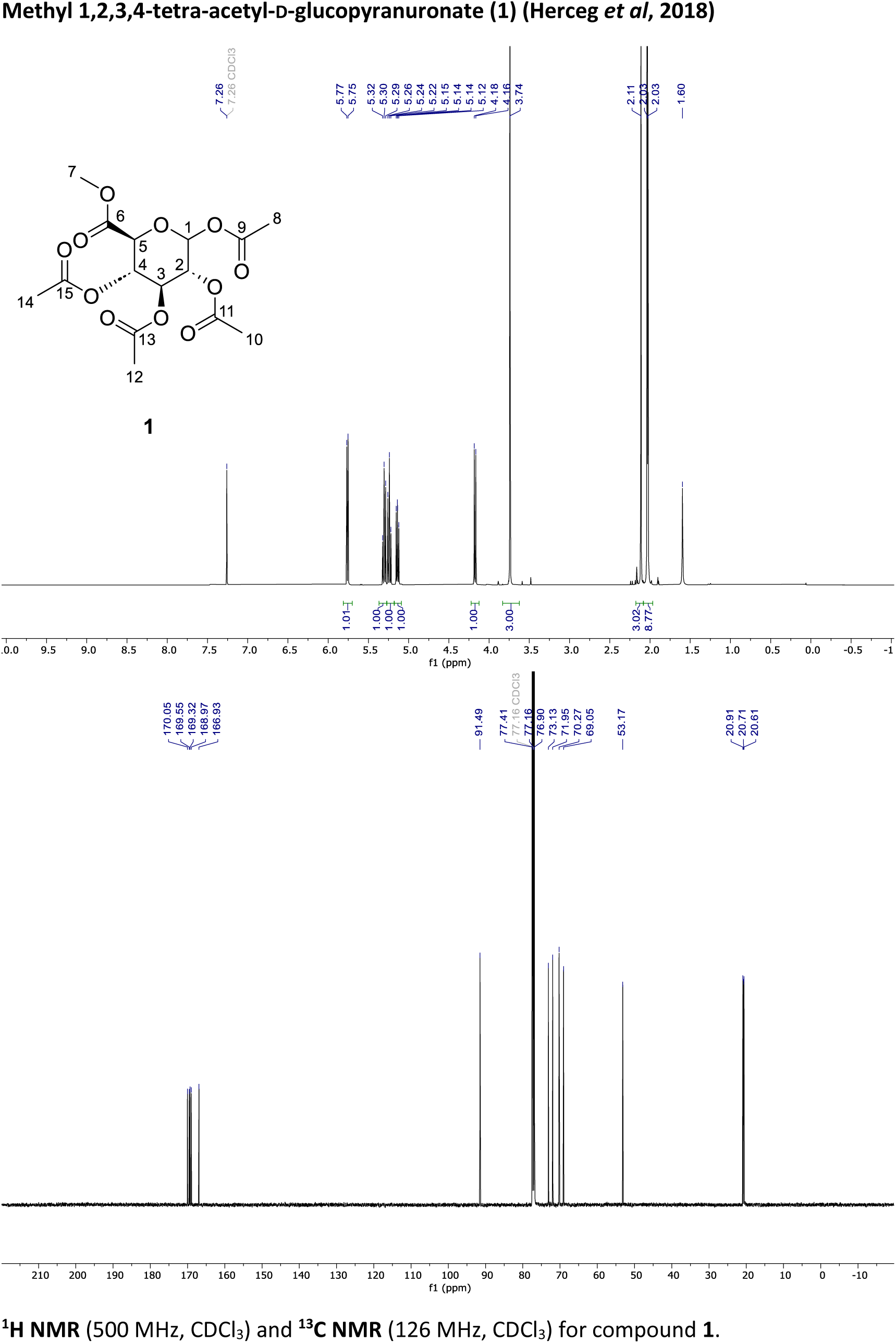

**^1^H NMR** (500 MHz, CDCl_3_) and **^13^C NMR** (126 MHz, CDCl_3_) for compound **1**.

### (2*R*,3*R*,4*S*,5*S*,6*S*)-2-Bromo-6-(methoxycarbonyl)tetrahydro-2*H*-pyran-3,4,5-triyl triacetate (2) (Walther *et al*, 2019)

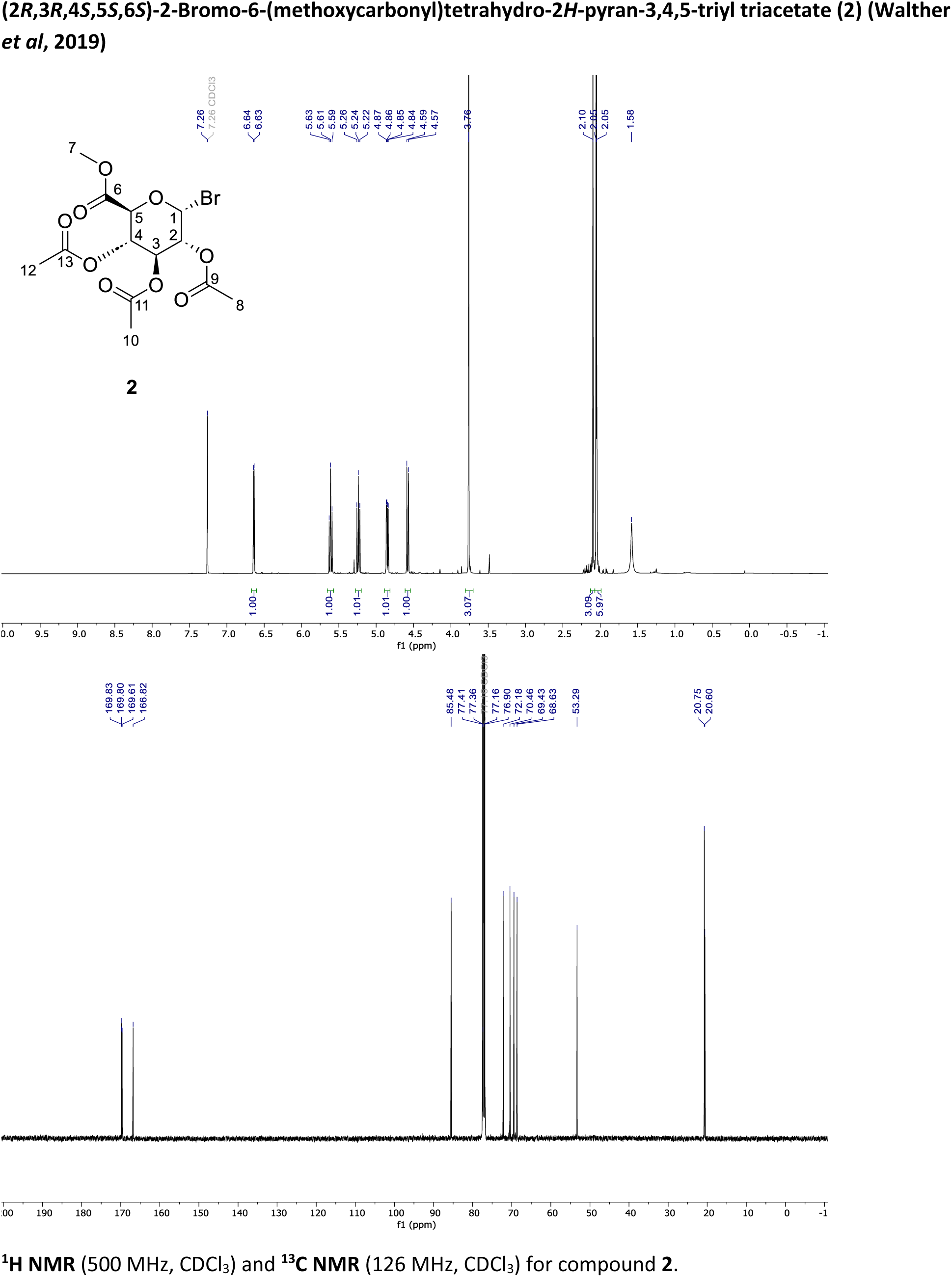

**^1^H NMR** (500 MHz, CDCl_3_) and **^13^C NMR** (126 MHz, CDCl_3_) for compound **2**.

### (2*S*,3*S*,4*S*,5*R*,6*S*)-2-(methoxycarbonyl)-6-((5-nitroquinolin-8-yl)oxy)tetrahydro-2*H*-pyran-3,4,5-triyl triacetate (3)

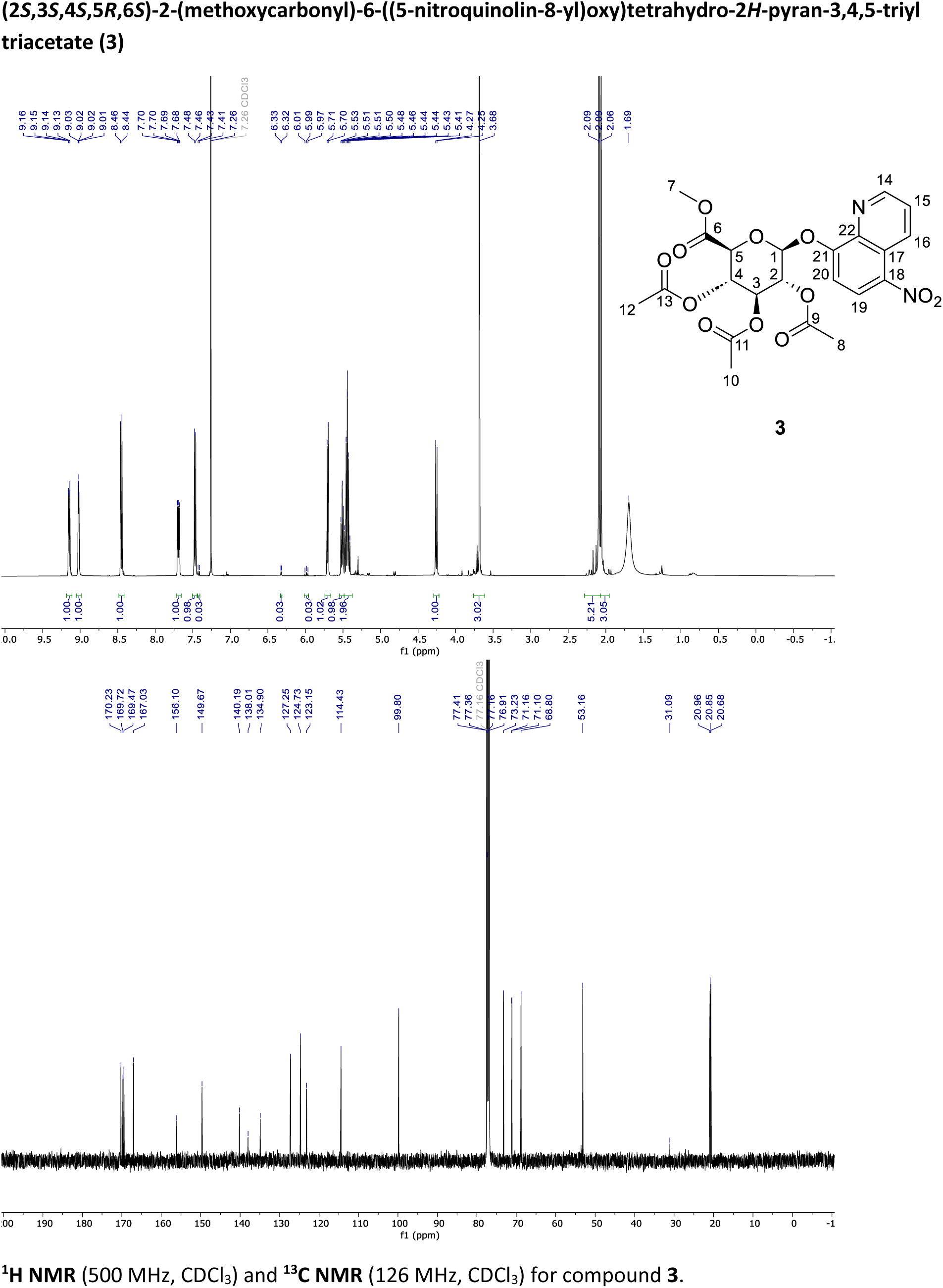

**^1^H NMR** (500 MHz, CDCl_3_) and **^13^C NMR** (126 MHz, CDCl_3_) for compound **3**.

### (2*S*,3*S*,4*S*,5*R*,6*S*)-3,4,5-Trihydroxy-6-((5-nitroquinolin-8-yl)oxy)tetrahydro-2*H*-pyran-2-carboxylic acid (4)

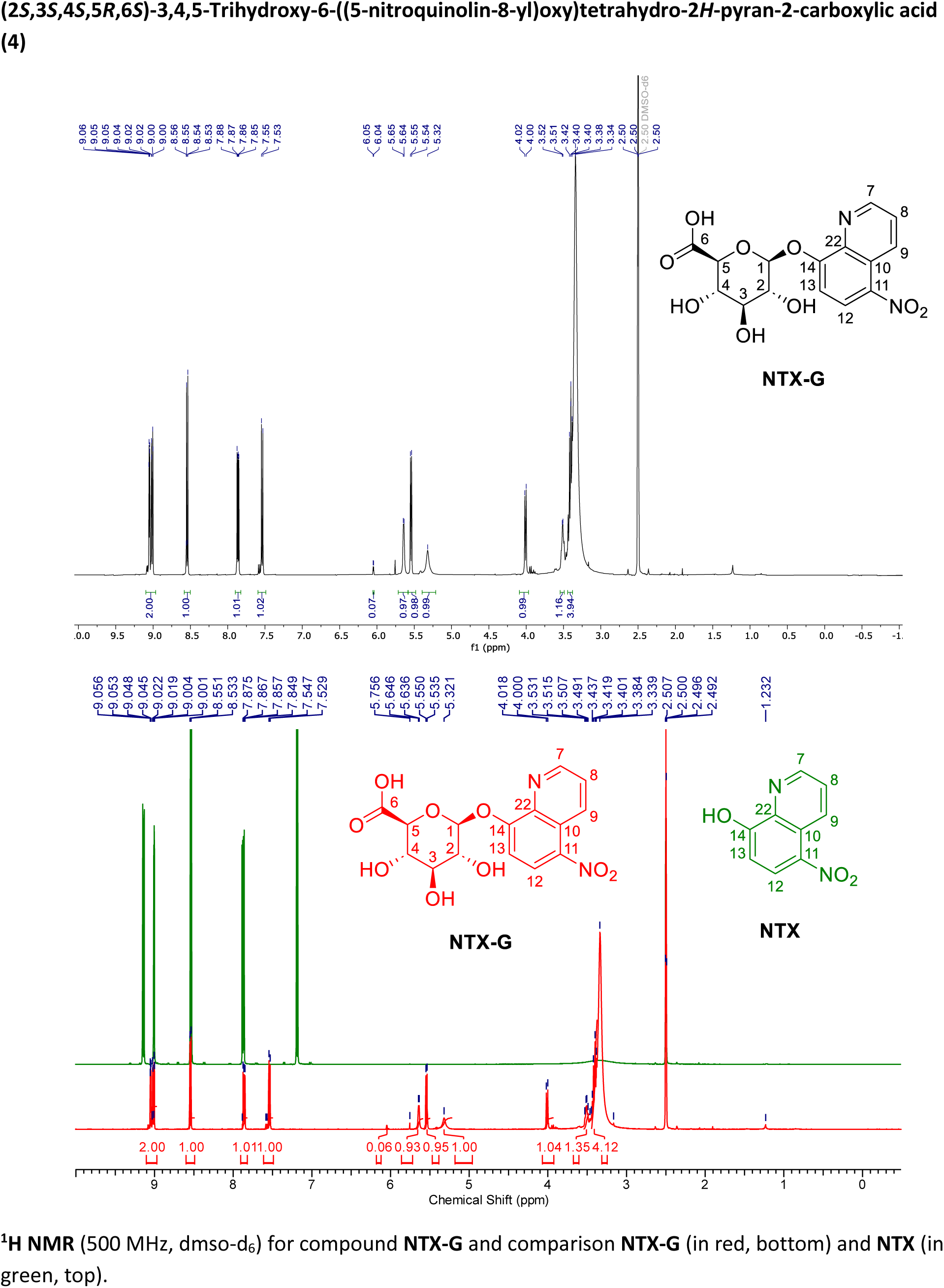

**^1^H NMR** (500 MHz, dmso-d_6_) for compound **NTX-G** and comparison **NTX-G** (in red, bottom) and **NTX** (in green, top).

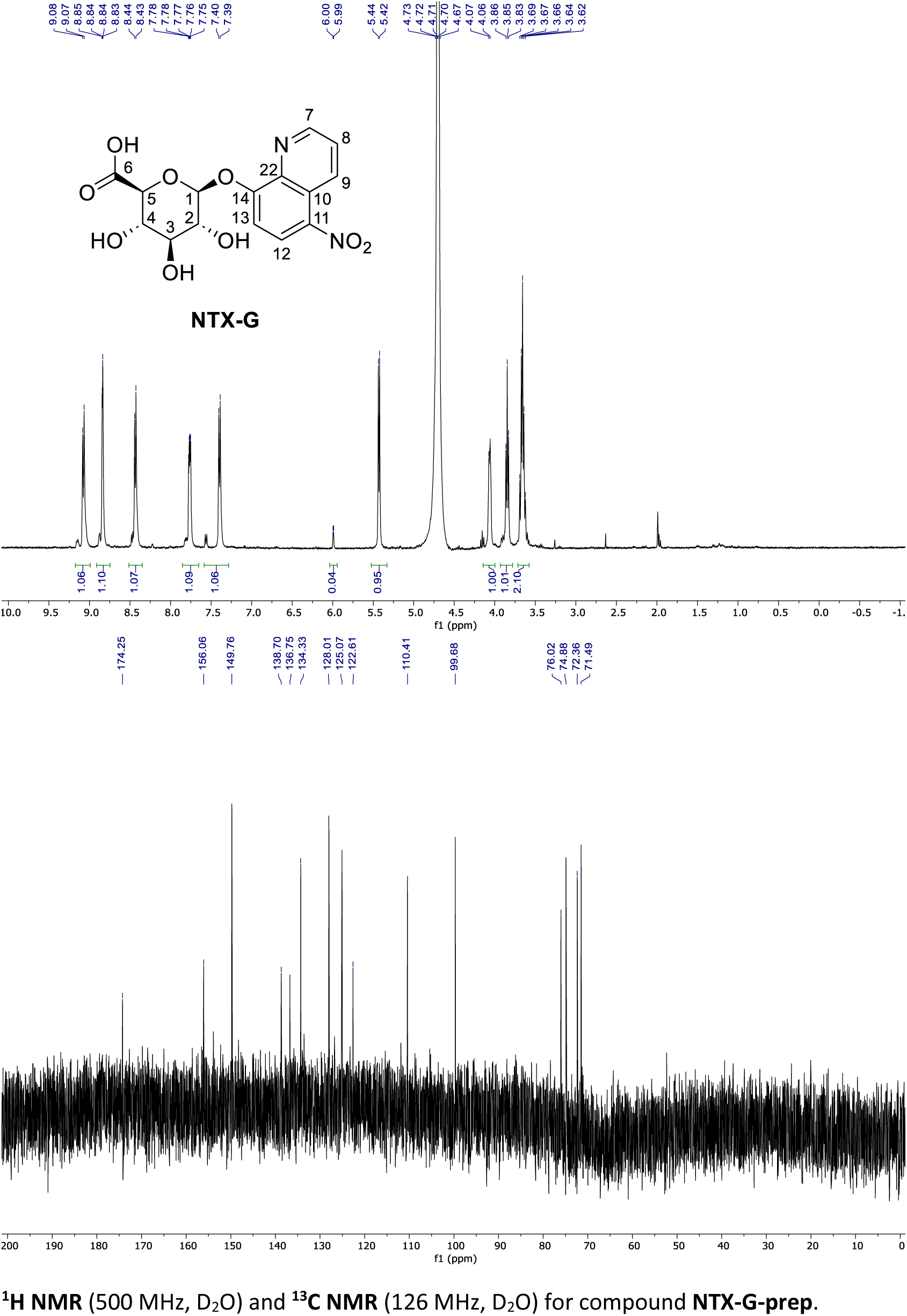

**^1^H NMR** (500 MHz, D_2_O) and **^13^C NMR** (126 MHz, D_2_O) for compound **NTX-G-prep**.

### 5-Nitroquinolin-8-yl hydrogen sulfate (NTX-S) (refer to Scheme SI 2)

A 2.6 mL dimethylacetamide solution of 5-nitroquinolin-8-ol (50.0 mg, 0.26 mmol) and SO3‧NEt3 complex (71.5 mg, 0.39 mmol) was heated in a crimp vail. After stirring for 10 min at 50 °C, the reaction mixture was slowly quenched with 10 mM HEPES buffer (pH 7.5) and directly purified via C18 revered-phase flash chromatography (gradient: MeCN/H_2_O, 0% MeCN to 10% MeCN) to yield NTX-S (54.0 mg, 0.20 mmol, 78%) as an amorphous white solid.

^1^H NMR (500 MHz, DMSO-*d*_6_) δ = 9.24 (d, *J* = 8.8 Hz, 1H), 9.05 (d, *J* = 3.6 Hz, 1H), 8.81 (bs, 1H), 8.57 (d, *J* = 8.8 Hz, 1H), 7.96 (dd, *J* = 8.7, 4.1 Hz, 1H), 7.24 (d, *J* = 8.8 Hz, 1H). ^13^C NMR (126 MHz, DMSO-*d*_6_) δ = 159.8, 148.7, 135.9, 135.2, 134.2, 129.5, 125.4, 122.8, 110.7. HRMS (ESI, *m/z*): calculated for C_9_H_7_N_2_O_6_S [M+H]^+^ 271.0019; 271.0019 found.

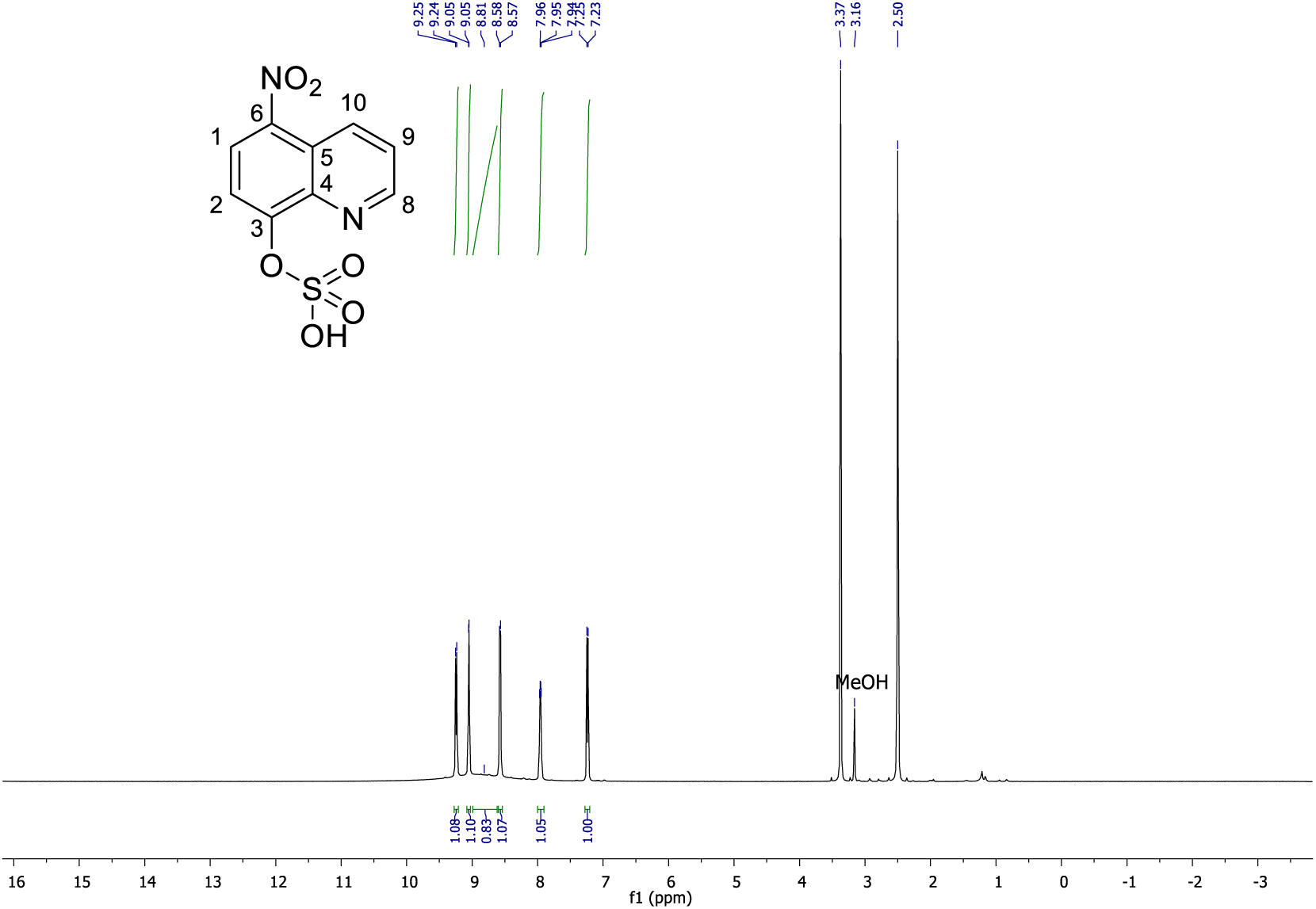

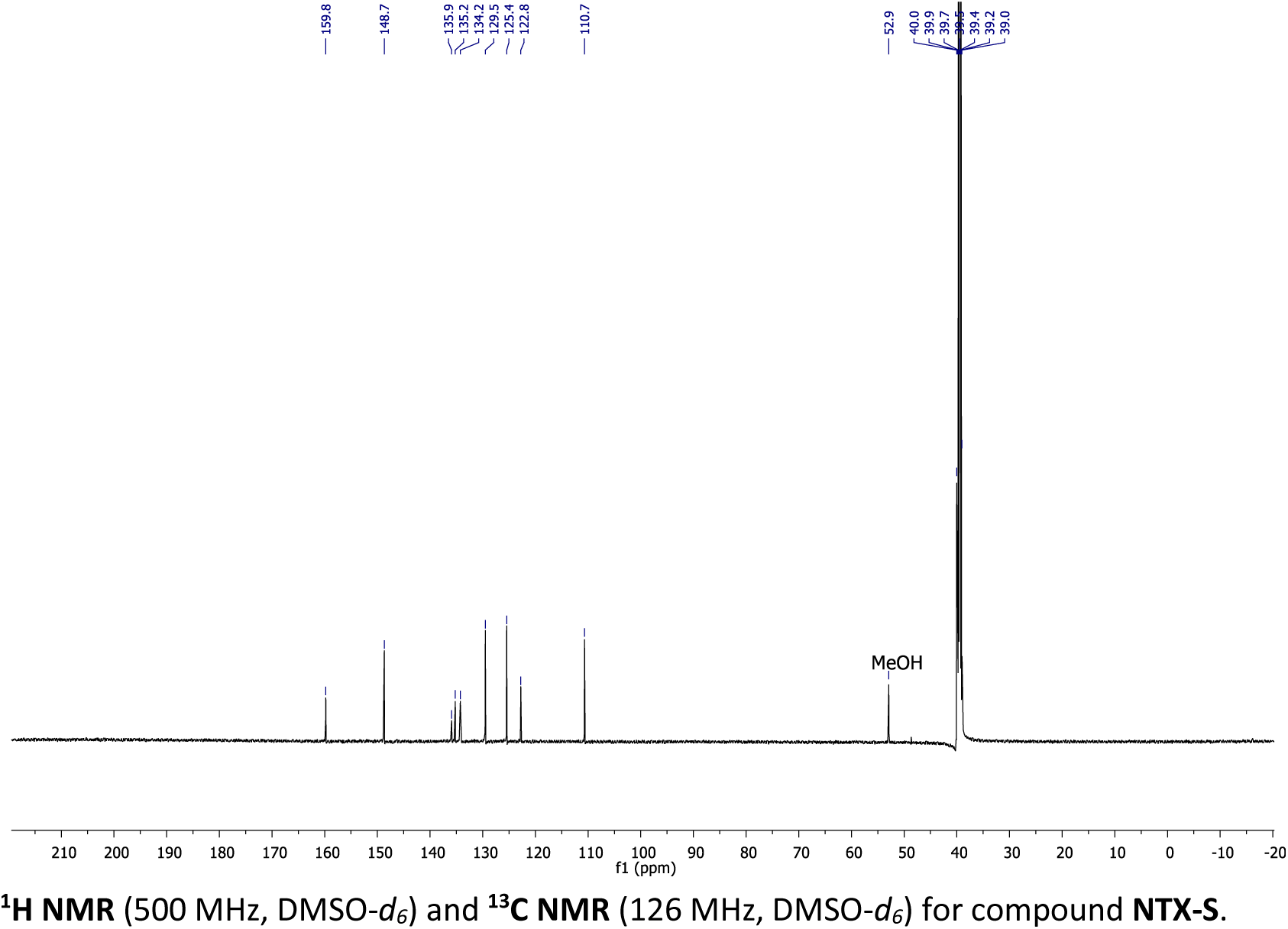

**^1^H NMR** (500 MHz, DMSO-*d_6_*) and **^13^C NMR** (126 MHz, DMSO-*d_6_*) for compound **NTX-S**.

